# The Biochemical Effects of Carotenoids in Orange Carrots on the Colonic Proteome in a Mouse Model of Diet-induced Obesity

**DOI:** 10.1101/2024.07.23.604335

**Authors:** Emilio Balbuena, Fadia Milhem, Buse Zeren Kiremitci, Taufika Islam Williams, Leonard Collins, Qingbo Shu, Abdulkerim Eroglu

## Abstract

Carotenoids are naturally occurring pigments in plants and are responsible for the orange, yellow, and red color of fruits and vegetables. Carrots are one of the primary dietary sources of carotenoids. The biological activities of carotenoids in higher organisms are well documented in most tissues but not the large intestine. The gastrointestinal barrier acts as a line of defense against the systemic invasion of pathogenic bacteria, especially at the colonic level. Proteins involved in tight junction assembly between epithelial cells and mucus secretion from goblet cells are essential for maintaining intestinal barrier homeostasis. A high-fat diet can cause gut impairment by inducing barrier permeability, leading to low-grade chronic inflammation via metabolic endotoxemia. Our hypothesis for this study is that the dietary intake of carotenoid-rich foods can alleviate obesity-associated gut inflammation and strengthen the intestinal barrier function. Male C57BL/6J mice were randomized to one of four experimental diets for 20 weeks (n = 20 animals/group): Low-fat diet (LFD, 10% calories from fat), high-fat diet (HFD, 45% calories from fat), HFD with white carrot powder (HFD + WC), or HFD with orange carrot powder (HFD + OC). Colon tissues were harvested to analyze the biochemical effects of carotenoids in carrots. The distal sections were subjected to isobaric labeling-based quantitative proteomics in which tryptic peptides were labeled with tandem mass tags, followed by fractionation and LC-MS/MS analysis in an Orbitrap Eclipse Tribrid instrument. High-performance liquid chromatography results revealed that the HFD+WC pellets were carotenoid-deficient, and the HFD+OC pellets contained high concentrations of provitamin A carotenoids, specifically α-carotene and β-carotene. As a result of the quantitative proteomics, a total of 4410 differentially expressed proteins were identified. Intestinal barrier-associated proteins were highly upregulated in the HFD+OC group, particularly mucin-2 (MUC-2). Upon closer investigation into mucosal activity, other proteins related to MUC-2 functionality and tight junction management were upregulated by the HFD+OC dietary intervention. Collectively, our findings suggest that carotenoid-rich foods can prevent high-fat diet-induced intestinal barrier disruption by promoting colonic mucus synthesis and secretion in mammalian organisms. Data are available via ProteomeXchange with identifier PXD054150.

## 1. Introduction

Carotenoids are naturally occurring pigments synthesized in plants, fungi, and some microorganisms (1). From a chemistry point of view, they are mainly C40 terpenoids (tetraterpenoids) and composed of eight isoprene units. While they are abundant in nature, only a few are found in our diet (2). Depending on their chemical structure, they can be classified as either carotenes or xanthophylls. Carotenes, such as α-carotene, β-carotene, and lycopene, are hydrocarbons, while their oxygenated derivatives are called xanthophylls, including β-cryptoxanthin, lutein and zeaxanthin (3). These are major dietary carotenoids detected in human serum (1). Among them, α-carotene, β-carotene, and β-cryptoxanthin can function as vitamin A precursors; thus, they are referred to as provitamin A carotenoids. In addition to that, carotenoids can exert essential biological functions due to their antioxidant and immunological properties, thereby serving as proxy biological markers of general population health (4).

Carotenoids are mainly absorbed in the small intestine in mammalian organisms (5). Cholesterol, fatty acids, phospholipids, monoacylglycerides, bile salts, and other hydrophobic molecules interact to produce mixed micelles with carotenoids. The bioaccessibility of individual carotenoids in naturally occurring mixed micelles is determined by their hydrophobicity, affecting the postprandial blood response (6). Once micelles form, absorption via transport receptors or passive diffusion occurs in the intestinal epithelial cells, depending on the dose (7). After micellization, carotenoids permeate through the intestinal mucus layer. This layer, structurally and characteristically determined predominantly by goblet and epithelial cells (8), is crucial for carotenoid absorption and serves as an immunological and physical barrier, regulating nutrient uptake and waste transit.

Carotenoids and their metabolites form chylomicron particles, enter the lymphatic system, and are absorbed for utilization in the systemic circulation (5). Factors such as bile acid salts and digestive enzymes may influence carotenoid transport through micelles (7). Reduced lipid droplet size can enhance mixed micelle formation and carotenoid incorporation, improving micellization. Still, excessive lipase activity may elevate free fatty acids, potentially lowering micelle pH, accelerating carotenoid degradation, and diminishing micellization efficacy (9). The transport of carotenoids from fruits and vegetables to the lipid phase of food, known as mass transfer, is significantly influenced by the food matrix, which primarily determines their bioavailability (10). β-Carotene from plants can have low bioavailability (10–65%) due to resistance from carotene-protein complexes, fibers, and plant cell walls during digestion, hindering adequate carotenoid release (11). Dietary β-carotene is mainly excreted through the large intestine due to poor bioavailability. The stability and bioavailability of carotenoids in the gut, the function of gastrointestinal bacteria in their metabolism, and the direct biological actions of carotenoids or their metabolites in the colon are among the critical knowledge gaps (12,13). Recent developments in proteomics have provided the most comprehensive view of the complex connections between carotenoid intake, gut health, and obesity, highlighting the need for additional integrative study in this area (1). Since the role of carotenoids in obesity-induced colon dysfunction is not established, we utilized a model of diet-induced obesity as carotenoids exhibit anti-obesogenic effects in other organs like the liver and adipose tissue (14,15).

Cellular uptake of carotenoids is facilitated by different molecules, including cluster determinant 36 (CD36) and scavenger receptor class B type 1 (SR-B1), that are highly expressed in proximal sections of mice intestines and less expressed in distal sections (16). Provitamin A carotenoids regulate intestinal protein expression for carotenoid absorption, with retinoids partly controlling SR-BI activity. Studies using mouse models and human cell lines show that dietary precursors produce retinoic acid via β-carotene 15,15’-dioxygenase (BCO1), inducing homeobox transcription factor intestine-specific homeobox (ISX) to suppress BCO1 and SR-B1 expression, impacting carotenoid conversion and uptake (17). Multiple dietary factors can influence the absorption of carotenoids. Fatty acids such as eicosapentaenoic acids increase SR-BI expression in Caco-2 cells (18). Moreover, it has been observed that dietary glucose leads to heightened expression of SR-BI in both Caco-2 cells, a widely used model for studying intestinal absorption, and in the intestines of mice, suggesting a potential regulatory role for glucose in modulating the absorption process of nutrients (19).

The presence of mucin and the charge of particles facilitate proper mucus adhesion, enabling mixed micelles to traverse the mucus layer and enhance the bioaccessibility of carotenoids (20). Mucins, particularly mucin-2, play a critical role in the mucus layer’s structure and function (21). For carotenoids, mixed micelles penetrate mucus to reach intestinal cells, releasing them to enter cells passively or via lipid transporters. Lipid transporters are crucial for carotenoid and lipid absorption, as lipid modulation can affect carotenoid uptake by altering transporter expression or interaction (20). Linoleic acid-rich glycolipids were found to enhance PPARγ and SR-B1 expression in rat intestines, improving fucoxanthin bioavailability (22), with further support from a Caco-2 cell study indicating increased SR-B1 mRNA expression in the presence of linoleic acid (23).

Metabolic syndrome is a disorder that increases the risk of type 2 diabetes and cardiovascular diseases as it is characterized by increased abdominal obesity, hyperglycemia, higher blood pressure, reduced HDL cholesterol, and hypertriglyceridemia (24). Furthermore, obesity due to Westernized high-fat diet (HFD) is associated with intestinal hyperpermeability via the promotion of intestinal inflammation and modulation of epithelial tight protein organization through signaling toll-like receptor (TLR) and nuclear factor kappa-light-chain-enhancer of activated B cells (NF-κB) signaling cascades (25). This HFD-induced proinflammatory response can lead to the development of intestinal pathologies like Crohn’s disease and ulcerative colitis, which are subsets of inflammatory bowel disease (IBD), as well as irritable bowel syndrome and celiac disease (25,26).

Carotenoids have a regulatory effect on multiple mediators related to inflammation and chronic diseases. These include C-reactive protein (CRP), proinflammatory cytokines (interleukin -1, -8, and -6), NF-κB, and tumor necrosis factor-alpha (TNF-α). Carotenoids can modulate the levels of these mediators by modulating oxidative stress or by promoting the overexpression of antioxidant and cytoprotective Phase II enzymes via the nuclear factor-erythroid 2-related factor 2 (Nrf2) and peroxisome proliferator-activated receptor (PPAR) pathways (27). Studies suggest ingesting carotenoids may help with obesity and associated concerns such as low-grade inflammation, insulin resistance, and hepatic steatosis (14). With an odds ratio of 0.66, a meta-analysis of 11 studies showed a negative association between total carotenoids and metabolic syndrome (28). The most significant inverse correlation was observed for β-carotene, followed by α-carotene and β-cryptoxanthin (28). A study demonstrated that treatment with fucoxanthin holds promise in preventing obesity by modulating intestinal homeostasis and alleviating gut dysbiosis and inflammation in a high-fat diet-induced mouse model. In male C57BL/6J mice fed a high-fat diet, 0.1% fucoxanthin supplementation from brown kelp *U. pinnatifida* reversed HFD-induced gut microbiota dysbiosis. This was achieved by inhibiting obesity- and inflammation-related *Erysipelotrichaceae* and *Lachnospiraceae* and encouraging the growth of beneficial bacteria such as *Bifidobacterium*, *Lactobacillus/Lactococcus*, and certain butyrate-producing bacteria (29). Additionally, fucoxanthin treatment increased butyrate levels in the colon contents of HFD-fed mice, improving insulin sensitivity and alleviating HFD-induced obesity. Consequently, fucoxanthin treatment could reverse HFD-induced gut microbiota dysbiosis (29).

The gut, which has a high lymphocyte population, is an essential barrier against toxins and antigens. Vitamin A is vital for preserving immunological homeostasis in this barrier, as its biologically active metabolite, retinoic acid, stimulates T and B cell expression of the gut-homing receptor, promotes regulatory T cell conversion, and causes T cell-independent IgA changes in naïve B cells (30). Intestinal immunity is weakened by vitamin A deficiency, increasing vulnerability to bacterial infections. Along with resistance to nematode infections, vitamin A-deficient mice have an increase in type 2 innate lymphoid cells that produce IL-13 (31). Other studies have shown that the altered immune profile in ISX-deficient mice may result from shifts in intestinal carotenoid processing. These mice show increased intestinal retinoic acid levels on a β-carotene-rich diet, synthesized in dendritic cells of lymphoid organs and in epithelial and stromal cells of mucosal lymphoid tissues where BCO1 is expressed (5). However, many of these studies have concentrated on administering individual compounds rather than whole foods, disregarding the effect of the pre-digestion process and focusing on a limited number of targets.

Advanced analytical techniques, particularly hyphenated approaches (e.g., LC-MS/MS) and multi-omics approaches are essential for identifying carotenoids and their metabolites along with their potential biological effects. Integrated analysis of proteomics, transcriptomics, metabolomics/lipidomics, and metagenomics is critical for improving our understanding of the bioactivity of natural components; carotenoids as a key focus with -omics studies has been highlighted by our team and collaborators (1). Proteomics explicitly allows studying the biological activities of carotenoid metabolites by detecting proteins that are differentially regulated within organs, tissues, or plasma, including subcellular fractions such as the cytosol, nucleus, or cell membrane. Unlike transcriptomics, it may better capture carotenoid bioactivity by accounting for posttranscriptional alterations and time-dependent mRNA expression, collecting additional biological variations (1). The usefulness of mass spectrometry-based proteomics as a method for determining the carotenoid status of populations was shown by our research team, Eroglu et al. (4), in which the metabolic effects of carotenoids concerning inflammation have been demonstrated in humans. In this study, we analyzed how dietary carotenoids in orange carrots modulated low-grade inflammation and impaired intestinal barrier function in obesity-associated colonic dysfunction.

## 2. Materials & Methods

### 2.1. Plant Material and Diet Formulation

Carrot varieties of two distinct colors (white and orange) were obtained from local supermarkets and then freeze-dried at the North Carolina Food Innovation Lab in Kannapolis, NC. All diet pellets were formulated at Research Diets, Inc. (New Brunswick, NJ, USA). Standard low-fat (D12450H, 10% calories from fat) and high-fat (D12451, 45% calories from fat) diet pellet compositions were selected; the sucrose content of D12450H and D12451 were matched. Lyophilized carrot powders were incorporated into the high-fat D12451 formula at 20% weight/weight (w/w). The dosage of 20% w/w has previously been utilized by our lab (32) and was selected to take account of the limited bioaccessibility of dietary carotenoids (33). Furthermore, rodents possess higher cleavage efficiency of carotenoids via BCO1 and ꞵ-carotene 9,10-dioxygenase (BCO2) than humans (15), so this dosage of 20% carrots is appropriate for the examination of intact, parent carotenoids rather than apocarotenoid cleavage products. Once formulated, all diets were irradiated (10 to 20 kGy) for sterility. The compositions of standard and custom diet pellets, as well as a vitamin mix (V10001C), have been provided in Supplementary Table 1.

### 2.2. Animals

Approval for the animal protocols in this study was obtained through the Institutional Animal Care and Use Committee (IACUC) at the North Carolina Research Campus. Male C57BL/6J mice (5 weeks of age, *n*=80) were purchased from The Jackson Laboratory (Bar Harbor, ME, USA). Two mice were co-housed per cage and fed a standard chow diet upon arrival in a temperature and humidity-controlled room with a 12-hour light/dark cycle. Following an acclimation period of seven days, mice were randomized into four dietary groups: (*n* = 20 per group): low-fat diet (LFD), high-fat diet (HFD), HFD with white carrot powder (HFD+WC), and HFD with orange carrot powder (HFD+OC). Since mice are coprophagic, fecal samples were grouped by the cage (*n*=10 per group for fecal endpoints). The dietary intervention lasted 20 weeks; water and experimental food pellets were administered *ad libitum*. Body weight for each mouse, the body weight change from baseline (before dietary intervention, week 0), and food consumption per cage were recorded every week. Following the 20-week dietary intervention, mice were anesthetized with isoflurane for cardiac blood and tissue harvest. Cardiac blood was collected in serum separator tubes (Sarstedt Inc., Newton, NC, USA) and placed at room temperature for 30 minutes, and serum was isolated by centrifuging the blood at 1,500 x g for 10 min at 4℃. Unexpelled feces still present in the colon were flushed out with saline. A portion of the distal colon was fixed in 10% formalin for histological examination. The remaining harvested tissue was snap-frozen in liquid nitrogen and then stored at −80℃ for further analysis.

### 2.3. Body Composition Analysis

Body composition measurements at baseline and week 20 of the study were analyzed with an EchoMRI-100 (EchoMRI, Houston, TX, USA). Mice were placed in a restraining cylinder without anesthesia and locked using the Velcro attachment. Total body composition (fat mass, lean mass, free water, and total water mass) was calculated by inserting the cylinder containing the mice into the chamber unit of the EchoMRI-100.

### 2.4. Biochemical assays

#### Serum Endotoxin and Lipopolysaccharide Binding Protein (LBP)

For endotoxin analysis, serum samples were diluted 1:5,000, and the manufacturer’s instructions were followed (PyroGene Recombinant Factor C Endpoint Fluorescent Assay-#50-658U, Lonza Bioscience, Walkersville, MD, USA). Serum samples were diluted at 1:50 for LBP analysis, conducted via the manufacturer’s instructions (Abcam, Waltham, MA, USA).

#### Serum C-Reactive Protein (CRP)

For CRP analysis, serum samples were diluted 1:2,000 (Mouse C-Reactive Protein/CRP Quantikine ELISA Kit #MCRP00, R&D Systems/Bio-Techne, Minneapolis, MN, USA) and the assay was executed following manufacturer’s instructions.

#### Fecal Calprotectin

Fecal protein was extracted via homogenization in an extraction buffer (0.1M Tris, 0.15M NaCl, 1.0M urea, 10 mM CaCl2, 0.1M citric acid monohydrate, 0.5 g/L BSA at a pH = 8.0) (34). Protein concentrations were measured with Rapid Gold BCA (Thermo Fisher Scientific, Waltham, MA, USA), and manufacturer’s kit instructions were followed for analysis using undiluted samples (Mouse S100A8/S100A9 Heterodimer DuoSet ELISA #DY8596-05 and DuoSet Ancillary Reagent Kit #DY008B, R&D Systems). Results were reported as calprotectin (pg)/total fecal protein (µg).

#### Colon IL-6

Distal colon tissues (approximately 30 mg) were homogenized in radioimmunoprecipitation assay (RIPA) buffer (Thermo Fisher) and incubated on ice for 30 minutes. Colon lysates were centrifuged at 16,000 xg for ten minutes, and the supernatants were collected. Protein quantification was conducted with Rapid Gold BCA, and manufacturer’s kit instructions were followed for analysis using undiluted samples (Mouse IL-6 DuoSet ELISA #DY406-05, R&D Systems).

### 2.5. Carotenoid Extraction and High-Performance Liquid Chromatography (HPLC)

Internal standards of all-trans-β-apo-8’-carotenal (purity ≥ 96%, catalog #10810, Sigma-Aldrich, St. Louis, MO, USA) for carotenoids and retinyl acetate (purity ≥ 95%, catalog #20242, Cayman Chemical, Ann Arbor, MI, USA) for vitamin A-related compounds were dissolved in HPLC-grade acetone and ethanol, respectively, and then added to all sample types before carotenoid extraction. Major dietary carotenoids and vitamin A-related compounds were used as external controls: α-carotene (purity ≥ 95%, catalog #19772, Cayman Chemical), β-carotene (purity ≥ 95%, catalog #16837, Cayman Chemical), and retinol (purity ≥ 95%, catalog #20241, Cayman Chemical). Dietary pellets (∼100 mg) were placed in the ZR Bashing Bead Lysis Tubes (Zymo Research, Irvine, CA, USA), and carotenoids were extracted via 500 μL HPLC-grade acetone followed by vibrant vortexing and centrifugation at 16,000 x g for five minutes at 4°C. The extraction steps were repeated two times, and supernatants were collected. Pooled supernatants were dried with nitrogen gas and reconstituted with 400 μL HPLC-grade acetone. Colon tissues (∼100 mg) were homogenized in 1 mL HPLC-grade hexane/acetone/ethanol (2:1:1 v/v/v) with internal standards and centrifuged at 16,000 x g for 10 minutes at 4°C. The step was repeated two times, and the supernatants were transferred to a fresh tube. Collected supernatants were centrifuged for the last time under the same conditions, followed by the nitrogen gas drying step, and reconstituted with 65 μL HPLC-grade acetone. Serum samples (60 μL) were placed in the Eppendorf tubes, and 180 μL HPLC-grade hexane/acetone/ethanol (2:1:1 v/v/v) was added to each sample. Centrifugation was done at 4,000 rpm for 1 minute at 4°C and repeated. The upper layer of the supernatant was collected, dried down with nitrogen gas, and reconstituted with 65 μL HPLC-grade acetone.

Extracted carotenoids from all sample types were detected by an Ultimate 3000 HPLC (Thermo Fisher Scientific). Separation of the carotenoids was achieved by an Acclaim C30 5 μm 4.6 x 150 mm column (Thermo Fisher Scientific), which was housed within a thermostatted TCC-3000 column oven (Thermo Fisher Scientific) held at 25°C. The instrument method involved a nine-minute isocratic equilibration period before sample injection-acetonitrile: methanol (v/v: 25:75 at time -9 to -0.5 min at a flow rate of 1.5 mL/min, 25:75 at time -0.5 to 0 min at a flow rate of 1.0 mL/min). The mobile phase was set as follows for the acquisition period at time zero-methyl tert-butyl ether: acetonitrile: methanol (v/v/v: 0:25:75 to 50:15:35 linear gradient at time 0–20 min, 50:15:35 isocratically at time 20–25 min) at a flow rate of 1.0 mL/min. The channels used for the detection of carotenoids and retinoids were as follows: all-trans-β-apo-8’-carotenal (464 nm), α-carotene (444 nm), β-carotene (450 nm), retinol (325 nm), and retinyl acetate (325 nm).

### 2.6. Quantitative Proteomics

#### 2.6.1. Sample Preparation for Quantitative Bottom-up Colon Proteomics

##### Extraction, Reduction, alkylation, digestion; Peptide clean-up

Peptide samples were prepared for proteomic analysis with the EasyPep Mini MS Sample Prep kit (#A40006, ThermoFisher). Whole-cell lysates were extracted from distal colonic tissue (approximately 30 mg; n = 4) using the EasyPep lysis buffer containing 1% nuclease. The Rapid Gold BCA kit determined the protein concentration according to manufacturer instructions, and samples were standardized to 100 μg protein with the EasyPep lysis buffer. Reduction and alkylation buffers were added to the standardized protein and incubated at 95°C for ten minutes on a heat block. Digestion into peptides was achieved via adding a trypsin/Lys-C protease mix and incubating with shaking for three hours at 37°C on a heat block, after which a stop solution was added to terminate the digestion. Hydrophilic/hydrophobic contaminants (i.e., buffer salts, detergents, and other biomolecules) were removed from the peptides by the clean-up columns from the EasyPep kit according to the manufacturer’s instructions. The resulting cleaned peptides were then dried by nitrogen gas.

##### Tandem Mass Tag pro (TMTpro) 16-plex Labeling

The dried peptides (100 μg) were reconstituted in 100 μL of 100 mM triethylamine bicarbonate (TEAB). TMTpro reagents (TMTpro 16-plex Label Reagent Set 0.5 mg #A44521, ThermoFisher) were dissolved in 20 μL of 100% LC/MS-grade anhydrous acetonitrile following equilibration to room temperature and the entire volume was added to the peptide samples of their respective treatment groups: LFD (126, 127N, 127C, 128N); HFD (128C, 129N, 129C, 130N); HFD+WC (130C, 131N, 131C, 132N), and HFD+OC (132C, 133N, 133C, 134N). Peptides were labeled for one hour at room temperature, after which the labeling reaction was quenched by incubation with 5 μL of 5% hydroxylamine for 15 minutes at room temperature. Equal amounts of each labeled peptide sample were combined into a single microfuge tube and dried down with nitrogen gas.

##### Peptide Fractionation

The dried, TMTpro-labeled peptides were dissolved in 300 μL of 0.1% trifluoroacetic acid solution and fractionated via high-pH, reversed-phase fractionation spin columns per the manufacturer’s instructions (Pierce™ High pH Reversed-Phase Peptide Fractionation Kit #84868, ThermoFisher). The fractions were eluted from the columns via an increasing gradient of acetonitrile and a decreasing gradient of triethylamine (Supplementary Table 2). Fraction 8 was dried with nitrogen gas and subjected to LC/MS analysis.

#### 2.6.2. nano-liquid chromatography-mass spectrometry (nanoLC-MS/MS)

The following LC-MS/MS experimentation was conducted by the Molecular Education, Technology, and Research Innovation Center (METRIC) at North Carolina State University. Materials were purchased from Fisher Scientific (Wilmington, DE): LC/MS grade water, LC/MS grade acetonitrile, LC/MS grade formic acid, PepMap™ Neo trap cartridge (C18, 5 μm particle, 300 μm × 5 mm). An Aurora Frontier analytical column (1.7 μm particle, 75 μm × 60 cm) was purchased from Ionopticks (Fitzroy, Victoria, AU).

Fractionated, TMTpro-labeled peptides were reconstituted in 100 µL water containing 2% acetonitrile and 0.1% formic acid. A 10 μL injection was analyzed by reversed-phase nano-liquid chromatography-mass spectrometry (nanoLC-MS/MS) using a Vanquish™ Neo UHPLC system (Thermo Scientific™, San Jose, CA, USA) interfaced with an Orbitrap Eclipse™ Tribrid (Thermo Scientific™) mass spectrometer. Peptides were concentrated, desalted, and separated using a trap and elute column configuration consisting of a PepMap Neo trap cartridge (Thermo Scientific™) in line with an Aurora Frontier analytical column held at 45°C. Mobile Phase A (MPA) consisted of water containing 0.1% formic acid, and Mobile Phase B (MPB) consisted of acetonitrile containing 20% water and 0.1% formic acid. The trap cartridge loading program used combined pressure (800 bar maximum) and flow control (100 μL/min maximum) with the loading volume automatically determined. For the analytical gradient, MPB increased from 2% at 0 min to 4% at 1 min, increased to 22% at 51 min, increased to 32% at 72 min, increased to 44% at 95 min, and increased to 50% at 105 min. The cartridge and column were washed for 10 min at 95% MPB. Mass spectrometer parameters were optimized for the detection and fragmentation of TMT-labeled peptides in a data-dependent experiment. Mass spectrometer parameters were set as follows: 1.8 kV positive ion mode spray voltage, ion transfer tube temperature of 275°C, master scan cycle time of 3 s, *m/z* scan range of 400 to 1,600 at 120 K resolution, standard AGC Target, automatic MS^1^ injection time, RF lens of 30%, intensity threshold for MS^2^ scan of 5.0e4, theoretical precursor fit threshold of 70% and *m/z* window of 0.7, dynamic exclusion for precursors applied for 90 s, mass resolving power of 50 K for data-dependent MS^2^ scans, *m/z* isolation window of 0.7, 38% normalized HCD collision energy, 200% normalized AGC Target, and 105 ms maximum injection time.

#### 2.6.3 Mass Spectrometry Data Analysis

Mass spectrometer raw data files were processed by Proteome Discoverer™ version 2.5 (PD2.5, Thermo Scientific) using a non-nested study design and a TMTpro quantification method. Within Chemical Modifications of PD, the TMTpro delta mass was defined as 304.207146 with possible modification at the protein N-terminus, any N-terminus, and possible residue targets H, K, S, and T. The TMTpro parameters were set according to specifications on the product data sheet for the TMTpro 16plex Isobaric Label Reagent Set, as shown in Supplementary Table 3. Reporter ion masses and their isotope correction factors were recorded in the Quan Channels dialog box. Residue modification of K and N-terminal modifications were selected. Mouse samples were categorized into 4 exposure groups: a low-fat diet, high-fat diet, high-fat diet white carrots, and high-fat diet orange carrots, and were associated with TMT quantification channels.

A *Mus musculus* protein database (92,954 sequences, taxonomy 10090) from Swiss-Prot and a custom home-built contaminants database containing human keratins and porcine trypsin were searched using PD2.5. Processing and consensus workflows were optimized for searching and quantifying TMTpro-labeled peptides. The Spectrum Files RC node used the following settings: *Mus musculus* protein database; trypsin (full) enzyme; 20 ppm precursor and 0.5 Da fragment mass tolerances; dynamic modification of TMTpro on any N-terminus; static modifications of TMTpro on K and carbamidomethyl on C; and non-linear regression with coarse tuning. The Spectrum Selector node used the following settings: MS1 precursor selection; isotope pattern re-evaluation; mass range from 350 – 5000 Da; FTMS mass analyzer; MS order of MS2; HCD activation; and scan type of full. The Reporter Ions Quantifier node used a 20 ppm integration tolerance and the most confident centroid integration method. The SEQUEST HT database search node used the following settings: *Mus musculus* and contaminants protein databases; trypsin (full) enzyme; maximum of 2 missed cleavage sites; minimum peptide length of 6 amino acids; 10 ppm precursor mass tolerance; 0.02 Da fragment mass tolerance; maximum of 3 equal dynamic modifications, which were N-terminal addition of acetyl group on methionine, oxidation or N-terminal loss of methionine, or loss of acetylated methionine; static modifications of carbamidomethyl on cysteine, TMTpro on any N-terminus, and TMTpro on lysine. Peptides were validated by the Percolator node with a *q*-value set to 0.05 and a strict false discovery rate (FDR) set to 0.01. The MSF Files node in the consensus step filtered peptide spectral matches with a maximum delta Cn of 0.05 and xCorr score of 1. The Reporter Ions Quantifier node used the following settings: use unique plus razor peptides; automatic reporter abundance, no quantitation value corrections; co-isolation threshold of 50; reporter signal-to-noise threshold of 10; SPS mass matches threshold of 65%; normalization using total peptide amount; scaling using “on controls average”; protein abundance based ratio calculations; no imputation; and ANOVA hypothesis testing.

### 2.7. Bioinformatic tools

#### Gene Ontology Network (GOnet) and STRING Protein-Protein Interaction Network

The GOnet maps for the HFD+OC vs. HFD and HFD+OC vs. HFD+WC comparisons were generated for functional enrichment analysis of proteins of interest by selecting job parameters: “Species: Mouse; GO namespace: biological process.” The GO term enrichment analysis type was chosen with a q-value <0.001. The output type of the analyzed results was an interactive graph with a *p*-value ≤ 9.70e-9 and formatted in the Cose layout selected with the unconnected annotation hidden. The software was accessed via GOnet using https://tools.dice-database.org/GOnet/ (35). The STRINGv12 software (https://string-db.org/) (36) was used for the protein-protein interaction network for the selected proteins. The protein names were listed under the “Multiple Proteins by Identifiers” tab, and the chosen organism was *Mus musculus*.

### 2.8. Immunohistochemistry

Distal portions of colon tissues were fixed in 10% formalin and washed with 70% ethanol. The tissue cassettes were then sent to the Anatomical Pathology Histology Lab in the College of Veterinary Medicine at North Carolina State University for processing. Once there, the colon portions were placed in a tissue processor for dehydration, clearing, and paraffin wax infiltration, followed by paraffin embedding and sectioning. Five-micrometer sections of colon tissues were stained with Alcian Blue or Mucin-2 (Abcam #ab272692), examined using a ZEISS Axio Observer microscopy platform (Carl Zeiss Microscopy, White Plains, NY, USA), and photographs were taken with AxioVision software. Briefly, Mucin-2 staining was achieved by a series of incubations: hydrogen peroxide (20 min), Fc receptor block (30 min), primary antibody MUC2 (1:4000 for 30 min), polymer rabbit secondary antibody (30 min), 3,3’-diaminobenzidine (DAB) chromogen (3 min), and hematoxylin counterstain (5 min). Mucin-2-positive (MUC2+) staining was quantified by ImageJ software (NIH, Bethesda, MD, USA) by converting the image to black and white 8-bit, and the default threshold of the grayscale was applied to the converted images. The percentage of area coverage by the stain (black) was measured via particle analysis through ImageJ’s “Analyze Particles” function. Unwanted staining (i.e., mucus masses between layers of tissue) was excluded from the analysis.

### 2.9. Statistical Analysis

Normality of distribution of the data was assessed via the D’Agostino-Pearson omnibus normality test for body weight, body composition, serum, and fecal analyses, while the Shapiro-Wilk test was used for colonic endpoints; *p* > 0.05 considered normally distributed. Equality of data variance was examined by using an F test with p > 0.05 considered equal variances. Then, one-way ANOVA and post hoc Tukey HSD were used to analyze the differences in endpoints across the various dietary intervention groups. The study employed a two-way mixed ANOVA with post hoc Tukey HSD to illustrate the differences in food consumption over time among multiple groups. A mixed-effect analysis using post hoc Tukey HSD was employed to examine the changes in body weight over time. The threshold for statistical significance was p < 0.05, and GraphPad Prism 9 was used for all statistical analysis (San Diego, CA, USA). The values within the text and figures were depicted as mean ± standard deviation (SD) or mean ± standard error of the mean (SEM) as indicated.

## 3. Results

### Orange Carrots are a Source of Provitamin A Carotenoids for Diet-induced Obese Mice

The content of dietary carotenoids (α-carotene, ꞵ-carotene) and vitamin A (retinol) was measured via HPLC in multiple specimens, including diet pellets, mouse serum, and mouse colon samples (**Table 1**).

**Table 1:** Content of dietary carotenoids (⍺-carotene, ꞵ-carotene) and vitamin A (retinol) measured via HPLC in **A)** diet pellets (*n*=3), **B)** mouse serum (*n*=10), and **C)** mouse colon (*n*=4) samples. ND: not detectable, below the limit of detection in HPLC. Values are means ± SD.

| Treatment groups | A) Diet Pellets ($\mu\text{mol/g}$) | | B) Serum ($\mu\text{M}$) | | | C) Colon ($\text{nmol/g}$) | | |
| --- | --- | --- | --- | --- | --- | --- | --- | --- |
| | α-carotene | β-carotene | α-carotene | β-carotene | Retinol | α-carotene | β-carotene | Retinol |
| LFD | 0.0 | 0.0 | 0.0 | 0.0 | $0.88 \pm 0.17$ | 0.0 | 0.0 | ND |
| HFD | 0.0 | 0.0 | 0.0 | 0.0 | $0.75 \pm 0.26$ | 0.0 | 0.0 | ND |
| HFD+WC | ND | ND | ND | ND | $0.92 \pm 0.11$ | ND | ND | ND |
| HFD+OC | $0.25 \pm 0.01$ | $0.18 \pm 0.01$ | $0.25 \pm 0.07$ | $0.26 \pm 0.06$ | $1.06 \pm 0.24$ | $0.72 \pm 0.28$ | $0.43 \pm 0.14$ | ND |

In the diet pellets, carotenoids were not detectable in the low-fat diet (LFD) and high-fat diet (HFD) formulations, indicating the absence or deficient levels of these compounds in these diets that could not be detected by the method used. Furthermore, the HFD supplemented with white carrot (HFD+WC) pellets also showed no detectable levels of carotenoids, suggesting that the white carrot component did not contribute significant amounts of α-carotene or β-carotene to the diet pellets. In contrast, the HFD supplemented with orange carrot (HFD+OC) pellets contained quantifiable amounts of carotenoids, α-carotene (0.25 ± 0.01 μmol/g) and β-carotene (0.18 ± 0.01 μmol/g). These results demonstrate how the variety of carrots consumed can significantly impact their carotenoid concentration. It’s also important to note that Research Diets, Inc. supplied 500,000 IU/g of vitamin A acetate, which was consistently incorporated into each treatment of diet pellets to achieve vitamin A-sufficient conditions.

The analysis of serum samples revealed findings that align with the carotenoid content observed in the diet pellets. There were no detectable carotenoids in the serum of mice fed the LFD, HFD, and HFD+WC, mirroring the absence of these compounds in the corresponding diet pellets. In mice fed the HFD+OC pellets, almost identical levels of α-carotene (0.25 ± 0.07 µM) and β-carotene (0.26 ± 0.06 µM) were measured in their serum. Furthermore, no significant variations were observed in the levels of circulatory retinol among any of the dietary groups, suggesting that vitamin A levels were constant irrespective of the diets. When examining the colon tissues of mice given the LFD, HFD, and HFD+WC, our findings showed that no noticeable concentrations of carotenoids were present. In contrast, colonic tissue extracts from mice fed HFD+OC had measurable amounts of α-carotene (0.72 ± 0.28 nmol/g) and β-carotene (0.43 ± 0.14 nmol/g). Despite the presence of these carotenoids in the HFD+OC group, retinol levels were undetectable in the colon tissues across all treatment groups.

### Carrot Supplementation on Body Composition and Metabolic Endotoxemia

Over time, analysis of body weight changes revealed distinct patterns among the dietary groups (**Figure 1A**). Starting at Week 2, all HFD groups exhibited significantly higher body weights than the LFD group. Between Week 9 and Week 14, mice in the HFD supplemented with white carrot and orange carrot groups had significantly higher body weights than those fed the HFD alone (*p* < 0.05). However, from Week 14 until the study endpoint at Week 20, only the body weight of the HFD+WC group remained significantly higher than that of the HFD group (*p* < 0.05). Conversely, as the experiment proceeded, the body weight of the HFD+OC group approximated that of the HFD and HFD+WC groups, suggesting an alignment in body weight among these groups.

**Figure 1:**
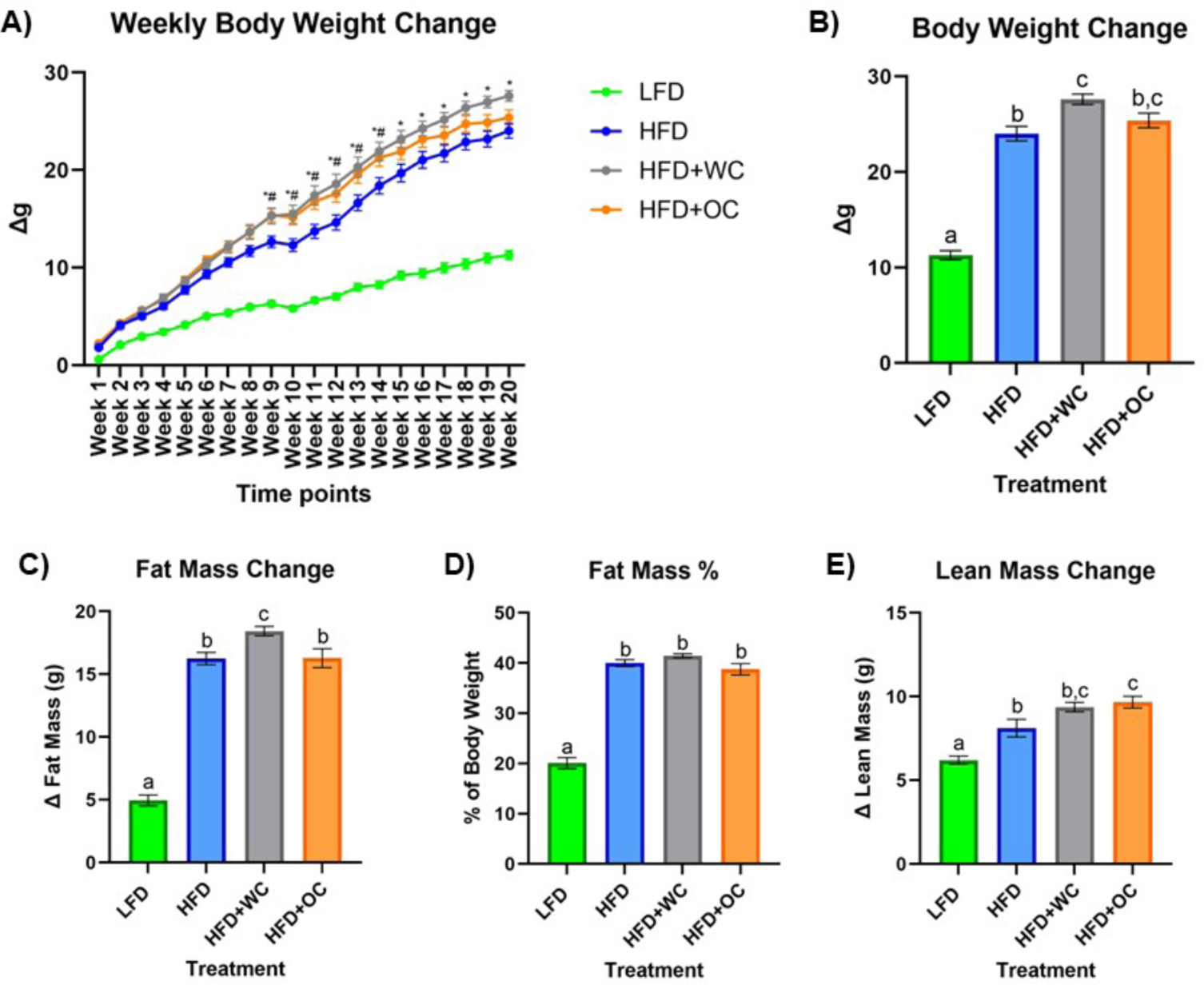
Dietary effects of carrots on HFD-induced body composition. **A)** Measurements of body weight gain on a weekly basis compared to baseline (week 0). *HFD vs HFD+WC p < 0.05, #HFD vs HFD+OC *p* < 0.05. **B)** Change in body weight at endpoint (week 20) compared to baseline. **C)** Change in fat mass determined by EchoMRI. **D)** Fat mass percentage of Week 20 body weight. **E)** Change in lean mass determined by EchoMRI. LFD, HFD, HFD+WC, HFD+OC n=20. Values are means ± SEM. Weekly body weight change was analyzed by two-way mixed ANOVA with post-hoc Tukey HSD. Changes in body weight, fat mass, fat mass percentage, and lean mass were analyzed by one-way ANOVA with post-hoc Tukey HSD. Letter differences indicate statistical significance (*p* < 0.05).

The body weight analysis of the different dietary groups provided clear distinctions (**Figure 1B**). Mice in the HFD group showed significantly higher body weights than those in the LFD (24.0 ± 0.8 g vs 11.3 ± 0.5 g, *p* < 0.0001), establishing the impact of a high-fat diet on body weight gain. Notably, mice in the HFD+WC treatment had significantly increased body weight than the HFD group (27.6 ± 0.5 g vs 24.0 ± 0.8 g, *p* < 0.001). In the comparison of the carrot-supplemented groups, no significant difference in body weight was observed between the HFD+WC and HFD+OC groups (27.6 ± 0.5 g vs. 25.4 ± 0.8 g), suggesting a similar influence of both types of carrot on body weight. Furthermore, the body weight of mice in the HFD+OC group was not significantly different from those in the HFD group (25.4 ± 0.8 g vs. 24.0 ± 0.8 g), suggesting that orange carrot supplementation did not add or reduce the weight gain compared to what was observed with the high-fat diet alone. These findings underscore the varying impacts of white and orange carrot supplementation on body weight under high-fat diet conditions.

At baseline, EchoMRI analysis revealed no significant differences in body weight across all treatment groups, indicating a consistent starting point for the study (**Supplementary Figure 1A**). However, after 20 weeks of treatment, fat mass gain measured by EchoMRI was significantly higher in all HFD groups, i.e., HFD (16.3 ± 0.5 g), HFD+WC (18.4 ± 0.4 g), and HFD+OC (16.3 ± 0.8 g), compared to the LFD group (4.9 ± 0.4 g, *p* < 0.0001) with HFD+WC consisting of the highest fat mass gain of all groups (**Figure 1C**). These findings demonstrate that all high-fat diet groups, irrespective of carrot supplementation, experienced a substantial increase in fat mass relative to the low-fat diet group throughout the study due to increased fat intake. Fat mass percentage of body weight at Week 20 is illustrated in **Figure 1D**, showing significantly lower fat mass in the LFD group (20.1 ± 1.1%) compared to all HFD treatment groups (*p* < 0.001), reflecting the influence of the assigned diets. Whereas the change in lean mass (**Figure 1E**) indicates that the HFD+OC group resulted in the highest lean mass gain (9.7 ± 0.3 g) among the groups with a significant increase compared to LFD (6.2 ± 0.2 g, *p* < 0.001) and HFD (8.1 ± 0.5 g, *p* < 0.05); no significant difference compared to HFD+WC lean mass (9.3 ± 0.3 g). Supplementary Figure 1 provides an extensive description of baseline and Week 20 values, which are essential for evaluating the effects of carotenoid intake throughout the study.

After 20 weeks on the specified diets, **Figure 2** depicts the levels of various targets as determined by ELISA in the serum, feces, and colon. The LFD group had significantly lower serum lipopolysaccharide (LPS) levels versus the HFD+WC group (*p* < 0.05), though there were no significant differences across the other groups (**Figure 2A)**. Lipopolysaccharide-binding protein (LBP) (**Figure 2B)** and C-reactive protein (CRP) (**Figure 2C)** demonstrated similar trends, with the LFD group having significantly lower levels of these markers in comparison to all other high-fat diet groups (*p* < 0.0001). These findings indicate that the low-fat diet is associated with lower systemic levels of inflammatory markers, though the carrot supplementation did not mitigate the HFD-induced gain. The amount of fecal calprotectin, a marker of intestinal inflammation (37), relative to total fecal protein was elevated in the HFD-fed animals. HFD+OC-fed mice showed significantly lower levels of fecal calprotectin compared to the HFD control (*p* < 0.05) (**Figure 2D)**. Similarly, the HFD+WC intervention also exhibited lower calprotectin levels than those on the HFD alone, though the reduction was more pronounced in the HFD+OC group. Additionally, the HFD+OC group showed the lowest concentrations of colonic interleukin-6 (IL-6) to total colonic protein (**Figure 2E)**; however, these variations were not statistically significant (*p* = 0.0664).

**Figure 2:**
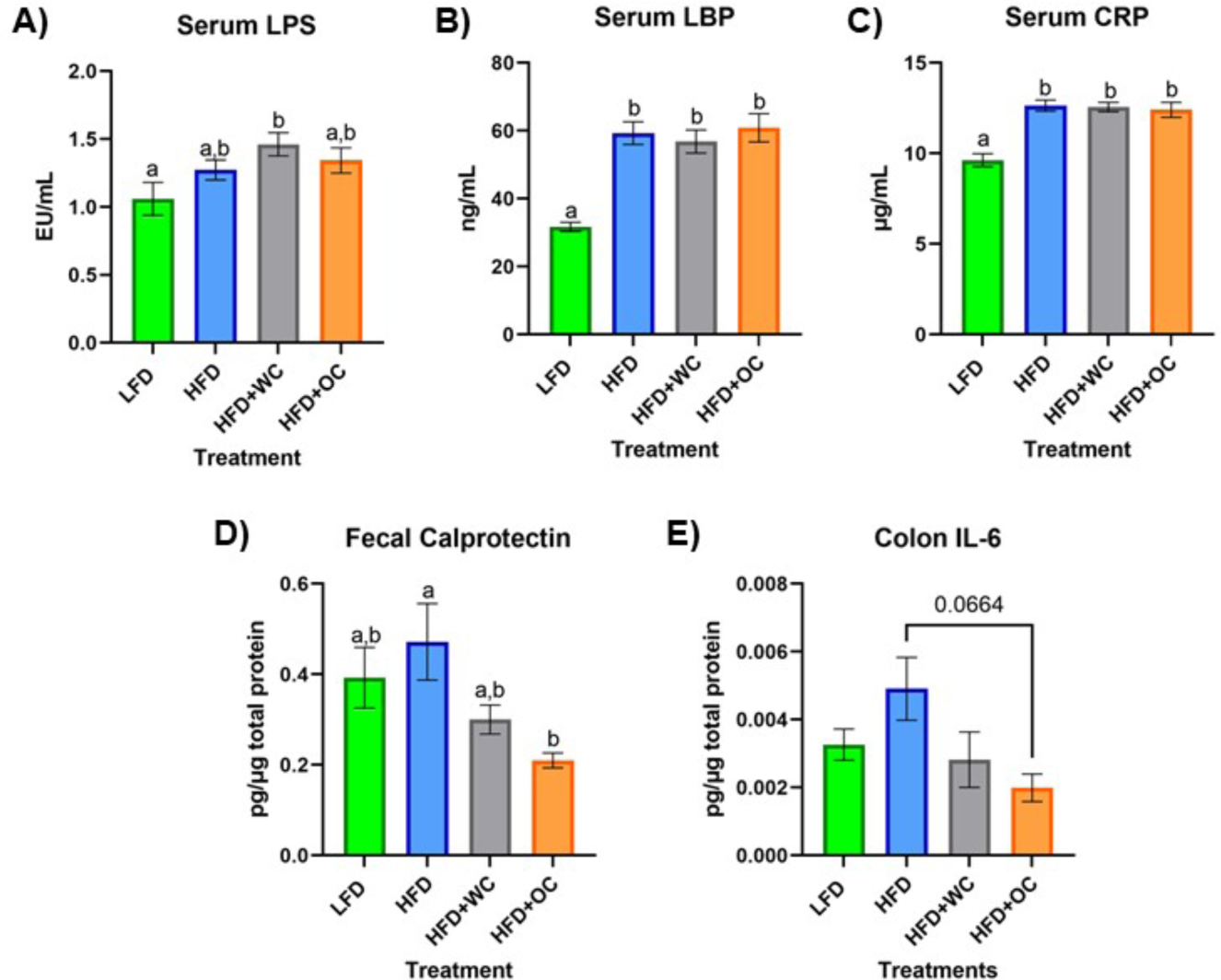
Serum, fecal, and colon ELISAs. **A-C)** Levels of circulating **A)** LPS, **B)** LBP, and **C)** CRP in the serum. **D)** amount of fecal calprotectin in total fecal protein. **E)** level of colonic IL-6 in total colonic protein. All analyses were determined via commercial ELISA kits. LFD, HFD, HFD+WC, HFD+OC n=10 for serum and fecal analyses; LFD, HFD, HFD+WC, HFD+OC n=4 for colonic analysis. Values are means ± SEM. ELISA measurements were analyzed by one-way ANOVA with post-hoc Tukey HSD. Letter differences indicate statistical significance (*p* < 0.05).

### Quantitative Proteomics: Key Comparisons between Dietary Intervention Groups

An outline of the quantitative, bottom-up proteomics process conducted on the distal colon tissue has been provided (**Figure 3A**). Volcano plots were generated to visualize differentially expressed peptides across two dietary intervention groups that significantly increased or decreased relative abundance. Surprisingly, there were no significantly regulated peptides in the comparison between LFD and HFD (**Figure 3B**). Supplementation of white carrots under high-fat conditions induced proteomic changes as seven peptides were significantly downregulated, and HFD+WC upregulated five peptides compared to HFD (**Figure 3C**). Furthermore, the downregulation of two peptides and the upregulation of ten peptides were induced in the HFD+WC group compared to LFD (**Supplementary Figure 3A**).

**Figure 3:**
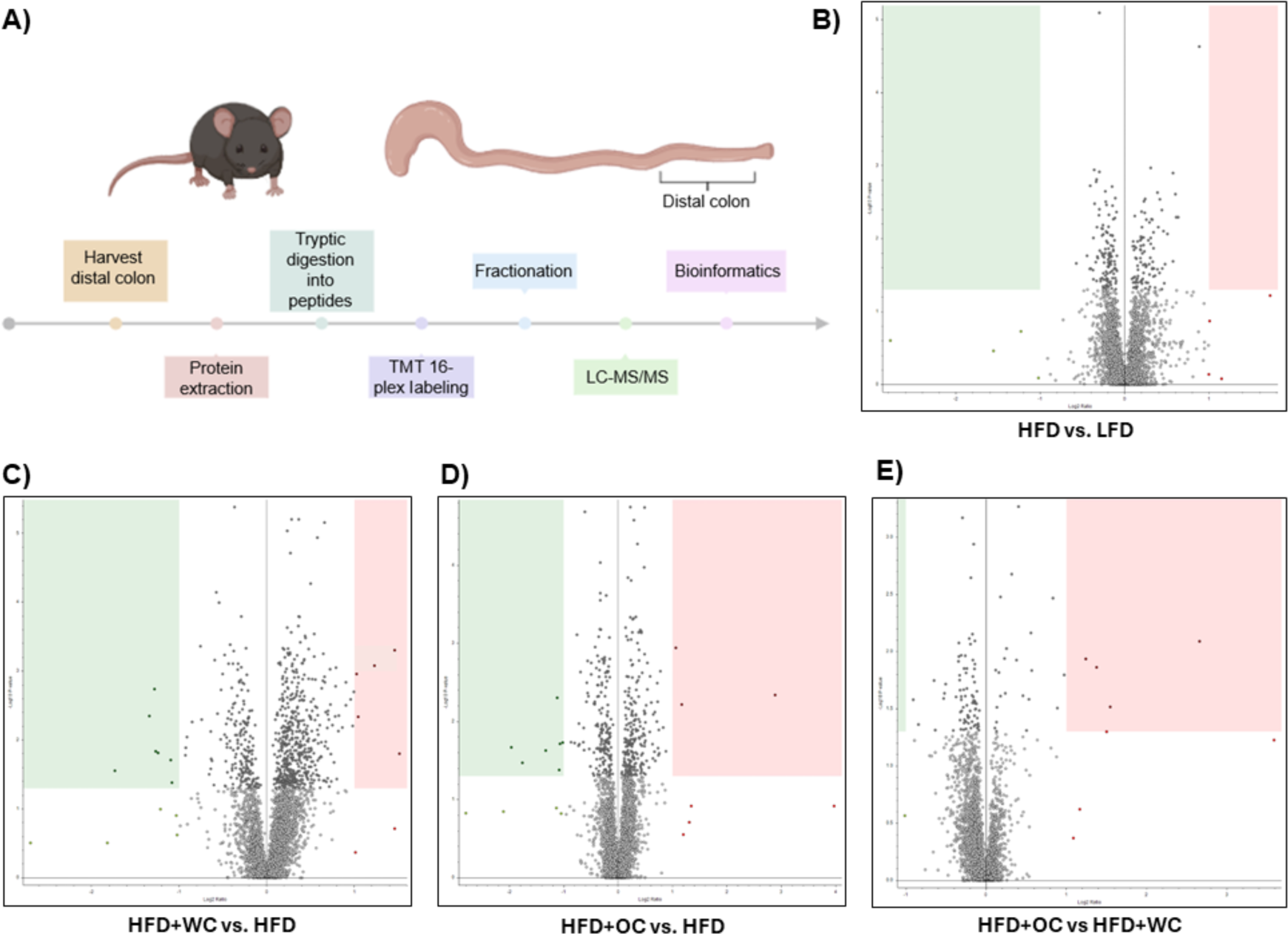
Quantitative proteomics on distal colon tissue. **A)** Outline of discovery bottom-up proteomics conducted with murine distal colon tissue. **B-E)** Volcano plots of differentially expressed peptides upon a comparison of abundances between **B)** HFD vs. LFD, **C)** HFD+WC vs. HFD, **D)** HFD+OC vs HFD, **E)** HFD+OC vs HFD+WC; boxes designate significant downregulation (green) or upregulation (red). LFD, HFD, HFD+WC, HFD+OC n=4 for colonic analysis.

Regarding orange carrots, seven peptides were significantly downregulated, three peptides were upregulated by HFD+OC compared to HFD (**Figure 3D**), and five peptides were upregulated versus LFD (**Supplementary Figure 3B**). Most notably, four peptides were significantly upregulated by the HFD+OC groups compared to the HFD+WC group (**Figure 3E**), indicating that carotenoids induce some proteomic changes. The Kyoto Encyclopedia of Genes and Genomes (KEGG) components of the proteins identified in the quantitative proteomics for Biological Processes, Molecular Function, and Cellular Components were detailed in **Supplementary Figure 3C-E**. The principal component analysis (PCA) plot depicted clustering for LFD and HFD+WC (**Supplementary Figure 3F)**. Less clustering across the four replicates was seen with the **HFD and HFD+OC**, as these two groups had a slight overlap.

### Mapping the Colon Proteome: HFD+OC versus HFD and HFD+WC

Gene Ontology Network (GOnet) maps have been generated for functional annotation and differential expression of proteins identified in the distal colon extracts. In particular, maps including the relative expression between HFD+OC versus HFD (**Figure 4**), HFD+OC versus HFD+WC (**Supplementary Figure 4**), and fold changes of various proteins have been provided (**Table 2**).

**Figure 4:**
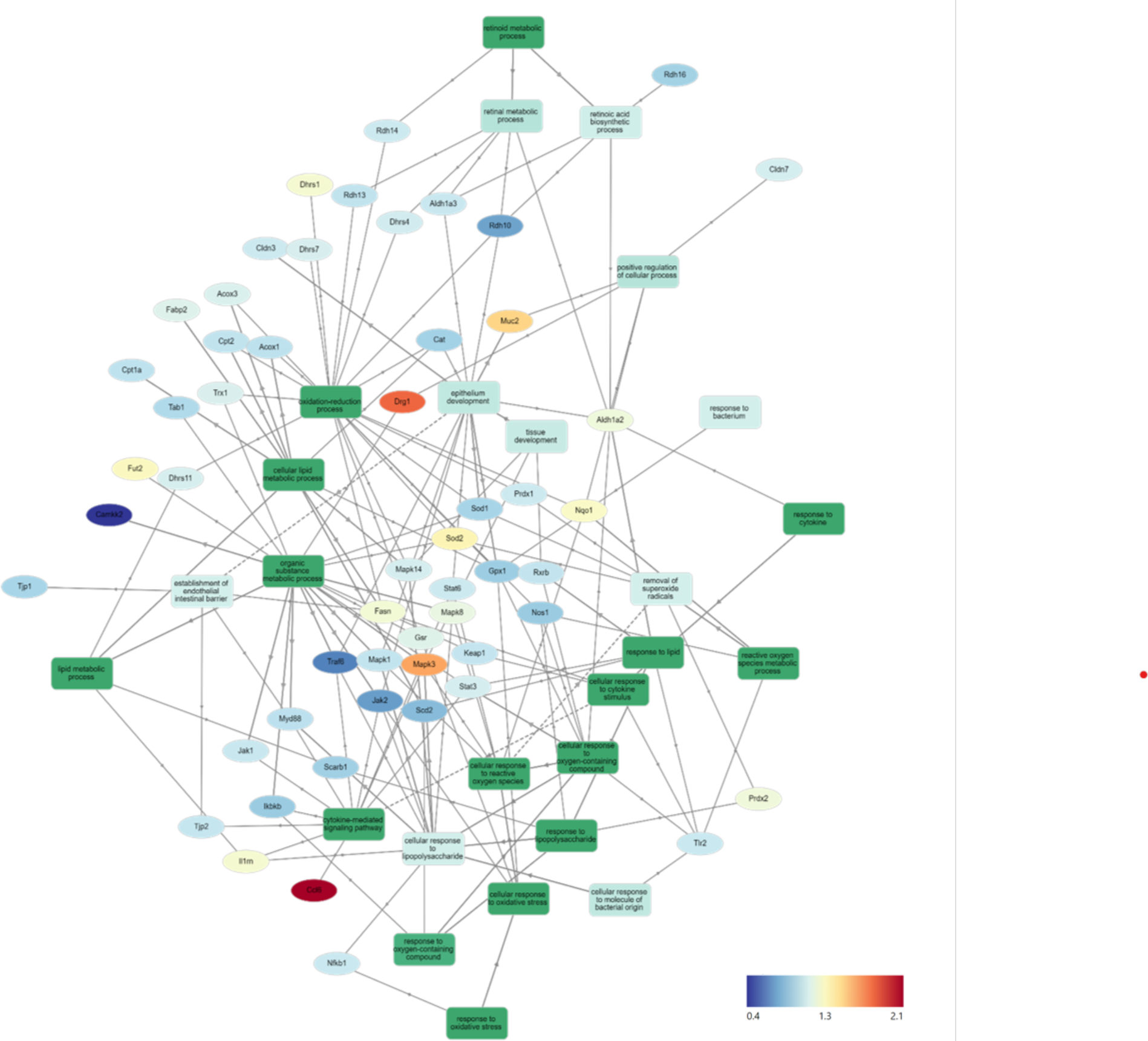
HFD+OC/HFD GOnet Map. Gene Ontology Network (GOnet) map of key differentially expressed proteins with a relative quantitative comparison between HFD+OC versus HFD (color of protein nodes and legend correspond to HFD+OC/HFD fold change). KEGG Pathway: Biological Process

**Table 2:**
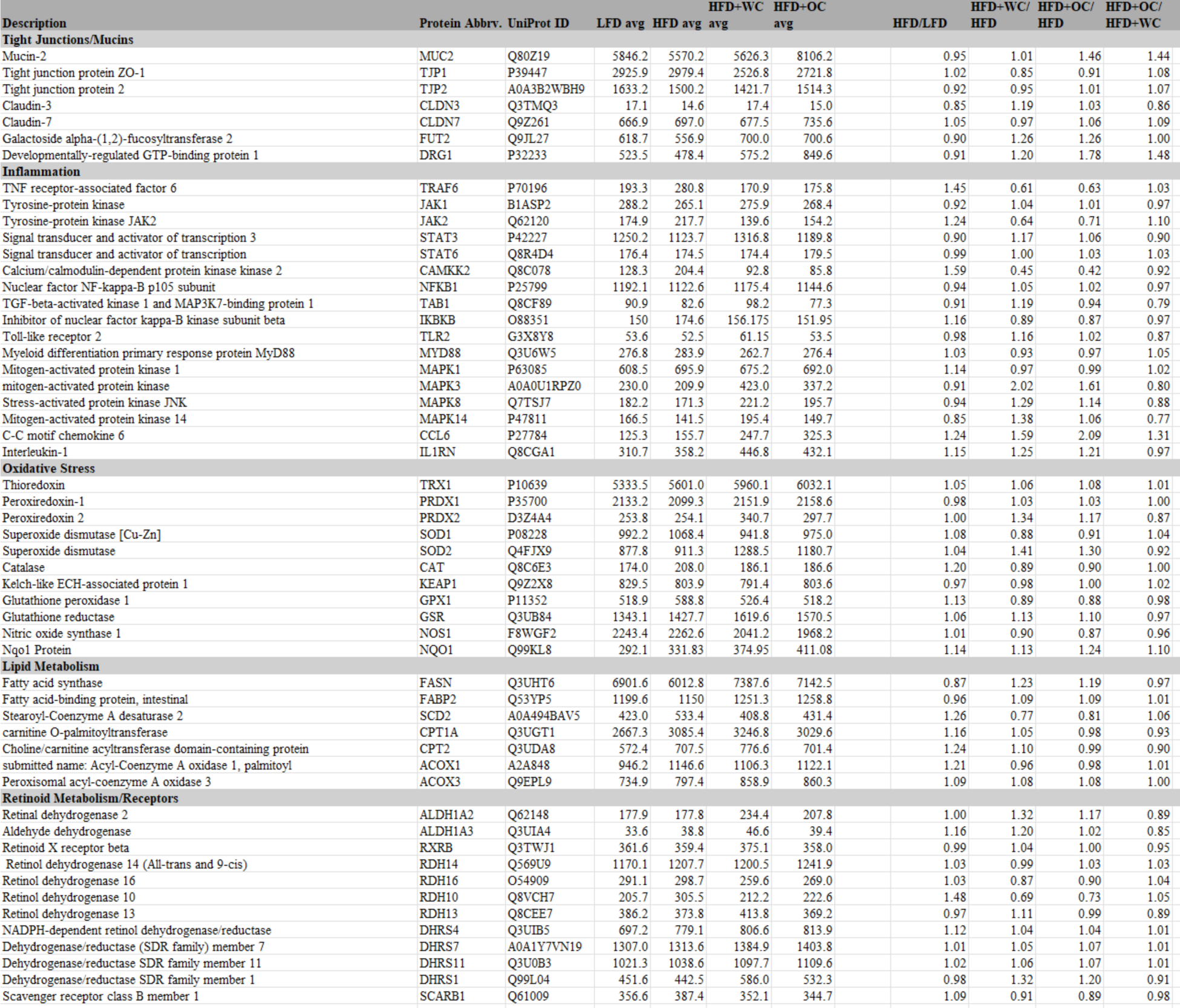
GOnet Fold Change. Average peptide abundances per treatment group and relative fold change comparisons of key identified proteins via Proteome Discoverer that were mapped in the GOnet plots of Figure 4 and Supplementary Figure 4.

Due to the documented anti-inflammatory and antioxidant efficacy of major dietary carotenoids, particular focus was placed on how the HFD+OC influenced the proteomic profile of significant players within these pathways. Notably, tumor necrosis factor receptor-associated factor 6 (TRAF6), an activator of NF-κB and mitogen-activated protein kinase (MAPK) signaling (38) (39), was significantly suppressed by both HFD+WC (*p* < 0.01) and HFD+OC (*p* < 0.01) compared to HFD abundance. However, there was no significant difference between the carrot groups. Unexpectedly, the abundance of toll-like receptor 2 (TLR2) and signaling partner myeloid differentiation primary response 88 (MYD88) (40) was not significantly affected by any dietary intervention groups, even when comparing LFD and HFD controls. Furthermore, players in the NF-κB signaling pathway such as the NF-κB p105 subunit (NFKB1), inhibitor of kinase NF-κB subunit beta (IKBKB/IKK-Β), transforming growth factor-β-activated kinase 1 regulatory subunit (TAB1), and interleukin 1 receptor antagonist (IL1RN) (38,41) did not differ in abundance. MAPK signaling was affected in various ways depending on the specific protein; for example, MAPK1 (also known as extracellular signal-regulated kinase 1, ERK2) was not significantly affected, while MAPK3 (also known as ERK1) was significantly upregulated by HFD+WC (*p* < 0.01) and HFD+OC (*p* < 0.05) relative to HFD. Furthermore, MAPK8 (also known as Jun N-terminal kinase, JNK) and MAPK14 (also known as p38) were significantly elevated by HFD+WC compared to HFD (*p* < 0.05) though no such changes were seen with HFD+OC.

Tyrosine-protein Janus Kinase 2 (JAK2) was also significantly downregulated by both HFD+WC (*p* < 0.05) and HFD+OC (*p* < 0.05) relative to HFD, along with its signaling partner signal transducer and activator of transcription 3 (STAT3) by HFD+WC (*p* < 0.01) and HFD+OC (*p* = 0.078); JAK1 and STAT6 were not affected by the treatment groups. C-C motif chemokine 6 (CCL6), which can act as an antibacterial peptide in the colonic mucosa (42), was highly upregulated by HFD+WC (*p* < 0.05) and to a greater extent by HFD+OC (*p* < 0.01) compared to HFD, also leading to a nearly significant difference between the carrot groups (*p* = 0.083). Fold change (FC) values of calcium/calmodulin-dependent protein kinase kinase 2 (CAMKK2), a promoter of tumorigenesis (43), were reduced in HFD+WC (fc = 0.45) and HFD+OC (fc = 0.56) versus HFD, though these changes did not reach significance due to sample variance.

Various proteins are involved in the cellular response to oxidative stress in the intestine (44,45), primarily regarding the breakdown of reactive oxygen species. However, several of the critical proteins involved in this antioxidant mechanism, including thioredoxin (TRX1), peroxiredoxin-1 (PRDX1), superoxide dismutase (SOD1), Kelch-like ECH associated protein (KEAP1), and glutathione peroxidase 1 (GPX1), NAD(P)H quinone dehydrogenase 1 (NQO1), and nitric oxide synthase (NOS1) were not influenced by any treatment groups. Catalase (CAT) was significantly increased by HFD alone compared to LFD (*p* < 0.05), as no other substantial changes were seen. Meanwhile, mitochondrial SOD2 was upregulated by HFD+WC compared to LFD (*p* < 0.05) and HFD (*p* < 0.05). Additionally, HFD+WC and HFD+OC led to an elevation of glutathione reductase (GSR) compared to LFD (*p* < 0.05, *p* < 0.05) but not to HFD. Most notably, PRDX2 was highly affected by the treatment groups as both HFD+WC and HFD+OC were significantly higher than HFD (*p* < 0.01, *p* < 0.05) and LFD (*p* < 0.01, *p* < 0.05) as well as significantly different from each other (*p* < 0.05) with a more substantial increase caused by HFD+WC of the two carrot groups.

Under the challenge of a high-fat diet, there was a chance that lipid metabolism-related proteins were affected at the colonic level (46); however, the only changes seen in these selected proteins were compared to the LFD group. β-Oxidation protein acyl-Coenzyme A oxidase 1 (ACOX1) was significantly higher in all HFD groups compared to LFD (*p* < 0.01, *p* < 0.05, *p* < 0.05) and ACOX3 was elevated in HFD+WC and HFD+OC relative to LFD (*p* < 0.05, *p* < 0.05). Furthermore, carnitine O-palmitoyltransferase (CPT1A) and choline/carnitine acyltransferase domain-containing protein (CPT2) were upregulated by HFD+WC compared to LFD (*p* < 0.01, *p* < 0.01). However, there were no significant changes in stearoyl-Coenzyme A desaturase 2 (SCD2) and intestinal fatty acid binding protein (FABP2). Fatty acid synthase (FASN), *de novo* lipogenesis of palmitate from malonyl-CoA and acetyl-CoA (47), was indeed upregulated by HFD+WC and HFD+OC compared to HFD (*p* < 0.01, *p* < 0.01).

Due to the supplementation of provitamin A carotenoids, enzymes involved in converting α-carotene and β-carotene into retinoids, whose roles have been detailed previously (5), were assessed. Scavenger receptor class B member 1 (SCARB1/SR-B1) was suppressed by HFD+OC compared to HFD (*p* < 0.05), which could indicate the negative feedback mechanism with ISX that down-regulates retinoid production from carotenoids under vitamin A-sufficient conditions (48). In line with this, proteins like retinal dehydrogenase 2 (ALDH1A2), aldehyde dehydrogenase 3 (ALDH1A3), retinol dehydrogenase 10 (RDH10), RDH14, RDH16, and dehydrogenase/reductase SDR family member 4 (DHRS4), DHRS7, DHRS11 were not significantly regulated by any treatment groups. Furthermore, the retinoid X receptor beta (RXRβ) transcription factor, which is activated by 9-cis-retinoic acid (49), was not significantly affected by any dietary intervention group. However, DHRS1 abundance was upregulated by HFD+WC and HFD+OC compared to HFD (*p* < 0.01, *p* < 0.05), and DHRS4 was elevated by HFD+WC and HFD+OC in relation to HFD (*p* < 0.01, *p* < 0.01).

Finally, proteins involved with intestinal epithelial homeostasis were highlighted due to apparent differential expression uniquely induced by the orange carrot group. Developmentally-regulated GTP-binding protein 1 (DRG1) was strongly elevated by HFD+OC compared to all groups, most notably when compared to HFD (*p* < 0.01) and also HFD+WC (*p* < 0.05). However, expression of colitis-associated galactoside alpha-(1,2)-fucosyltransferase 2 (Fut2) (50) was not significantly affected by any treatment group. Mucin-2 (MUC2), associated with epithelium development and tight junction proteins, was one of the highly upregulated proteins induced by HFD+OC supplementation compared to HFD (*p* < 0.01), which was also seen when compared to HFD+WC (*p* < 0.01). As this pathway appears to be significantly regulated by HFD+OC, we focused on proteins involved in colonic mucus activity, which will be elaborated on in the following section.

### Orange Carrot Feeding Upregulated Colonic Mucus Synthesis and Secretion

Proteins involved in mucus production and secretion were highly differentially expressed within the distal colon samples, with notable upregulation by HFD+OC feeding. For mucin-2 (MUC2), which is a major component of the protective mucus layer of the intestine (51), there was a significant increase in protein abundance in the HFD+OC group compared to all other groups (*p* < 0.01), there were no differences between the LFD, HFD, and HFD+WC groups (**Figure 5A**). The abundances of IgGFC-binding protein (FCGBP), a coreceptor of MUC2 (51), in both HFD and HFD+WC, were trending lower compared to LFD, though not statistically significant (**Figure 5B**). HFD+OC elevated FCGBP abundance to near substantial levels compared to the HFD and HFD+WC groups (*p =* 0.061, *p* = 0.054) and to a level higher than LFD, though insignificant. A similar trend was seen with calcium-activated chloride channel regulator 1 (CLCA1), a major facilitator of goblet cell secretion of MUC2 (52), since the abundance in the HFD+OC group was higher than both the HFD (*p =* 0.075) and HFD+WC groups (*p* < 0.05), though only significantly with the latter group (**Figure 5C**). The dietary intervention did not lead to statistical differences in abundances of zymogen granule membrane protein 16 (ZG16), an antimicrobial component of the mucus complex (53) (**Figure 5D**). Anterior gradient protein 2 homolog (AGR2) is involved in MUC2 assembly and folding (54); however, there were no significant differences in the relative abundance of this protein across the treatment groups (**Figure 5E**). Kallikrein-1 (KLK1), commonly associated with MUC2 though the exact function is unclear (53), was significantly upregulated by HFD+OC treatment compared to the LFD (*p* < 0.01) and HFD (*p* < 0.05) groups (**Figure 5F**). HFD+WC intervention also increased KLK1 compared to LFD (*p* < 0.05).

**Figure 5:**
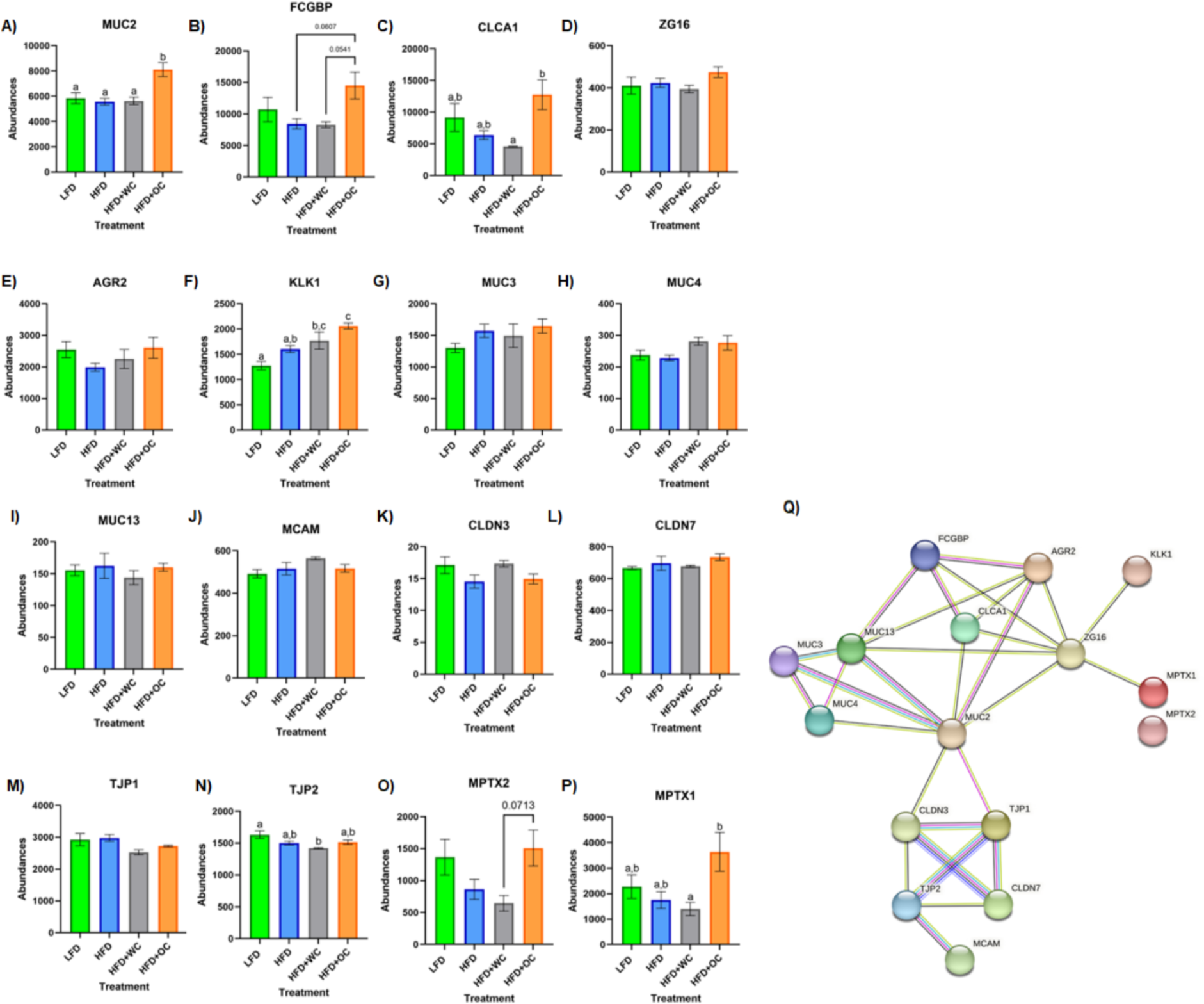
Upregulation of colonic mucus-associated proteins by HFD+OC. **A-P)** Quantitation for proteomic abundances of **A)** MUC2, **B)** FCGBP, **C)** CLCA1, **D)** ZG16, **E)** AGR2, **F)** KLK1, **G)**, MUC3, **H)** MUC4, **I)** MUC13, **J)** MCAM, **K)** CLDN3, **L)** CLDN7, **M)** TJP1, **N)** TJP2, **O)** MPTX1, **P)** MPTX2 identified via Proteome Discoverer. **Q)** STRINGv12 network map of selected mucus-associated proteins; experimentally determined (pink), curated databases (light blue), gene co-occurrence (dark blue), co-expression (black), text mining (mentioned together in Pubmed abstracts; yellow), protein homology (purple).

Other mucins were identified in the distal colon samples, such as mucin 3 (MUC3), mucin 4 (MUC4), mucin 13 (MUC13), and mucin 18 (MCAM); however, there were no significant differences among them (**Figure 5G-J**). Tight junction proteins were also identified in the colon tissue, like claudin-3 (CLDN3), claudin-7 (CLDN7), zonula occludens-1 (TJP1), and zonula occludens-2 (TJP2). No significant differences were observed in CLDN3 (**Figure 5K**) or CLDN7 (**Figure 5L**). Additionally, there were no significant differences in TJP1 (**Figure 5M)** and TJP2 (**Figure 5N**) except for a reduction of TJP2 in HFD+WC compared to LFD (p < 0.01). Another notable group of proteins significantly upregulated by orange carrot supplementation is the mucosal pentraxin (MPTX) family, which is involved in immune cell recruitment to the intestinal barrier (55). HFD+OC promoted the abundances of MPTX1 (**Figure 5O**) abundances compared to HFD (*p* = 0.078) and HFD+WC (*p* < 0.05) as well as of MPTX2 (**Figure 5P)** in comparison to HFD+WC (*p* = 0.071). STRINGv12 was utilized to demonstrate how these proteins were closely associated with each other, especially with MUC2 (**Figure 5Q**).

To obtain biochemical qualitative analyses of colon tissue samples, we assessed changes in colon morphology. Distal colon sections were paraffinized and treated with Alcian Blue to stain overall mucins in goblet cells (**Figure 6A**). While there appears to be a greater population of colonic goblet cells in the HFD+WC and HFD+OC groups compared to the LFD and HFD groups, there does not appear to be much difference across the carrot-supplemented groups. These distal colon sections were also stained MUC2 (**Figure 6B**), which indicated more differential expression of this specific protein. All HFD groups exhibited higher MUC2+ expression than the LFD group, and both carrot groups had more expression than the HFD group, reaching significance only by the HFD+OC treatment (*p* < 0.01). Though extensive MUC2+ staining is present in the colon tissue for both carrot groups, the staining intensity is higher in HFD+OC than in HFD+WC (*p* < 0.05).

**Figure 6:**
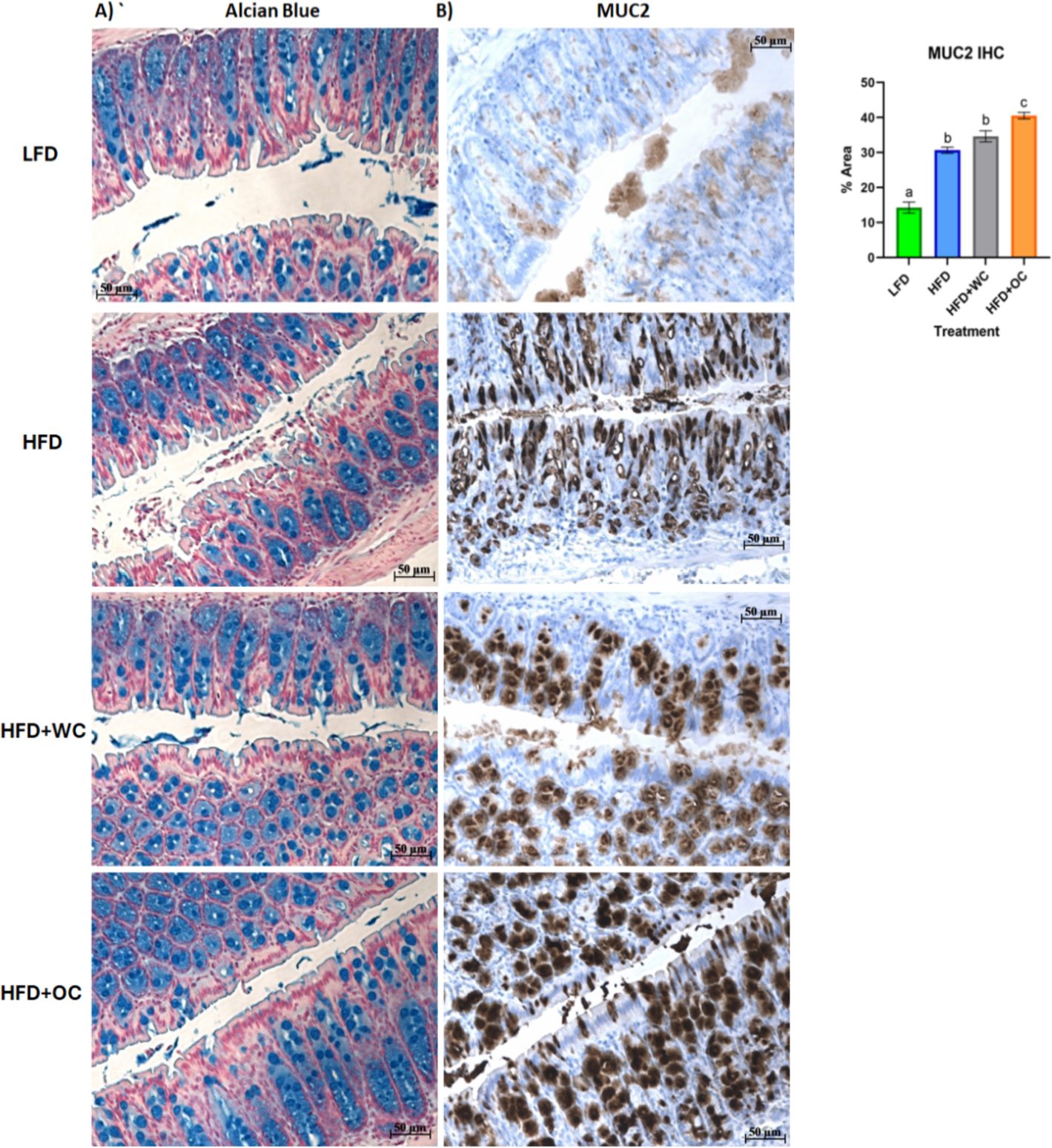
Distal colon histology of mucus-associated targets. **A)** Alcian Blue and **B)** Mucin-2 IHC staining. Scale bar of 50 µm included.

## 4. Discussion

Our objective in this study is to unravel the biochemical activities of major dietary carotenoids that affect obesity-induced metabolic syndrome, specifically in the context of the large intestine. We hypothesize that foods high in carotenoid content (i.e., orange carrots) could promote intestinal barrier function and reduce gut inflammation linked to obesity. Our results uphold this hypothesis by highlighting the potential advantages of carotenoids in promoting mucosal barrier integrity. Previously, our group assessed the levels of toll-like receptor 4 (TLR4), its co-receptor CD14, and the NF-κB p65 subunit to investigate the role of β-carotene in inhibiting the lipopolysaccharide (LPS)/TLR4 signaling pathway in human adenocarcinoma colon epithelial (HT-29) cells (56). Furthermore, we also looked at whether β-carotene modulated the levels of tight junction proteins occludin (OCLN) and claudin-1 (CLDN1) in that study and discovered that β-carotene decreased intestinal inflammation caused by LPS and increased tight junction protein levels, possibly improving barrier function (56).

We analyzed whether carotenoids derived from whole foods like orange carrots could reduce colon inflammation and impact host metabolism. In this study, 5-week-old male mice (n=20 per treatment) were fed an HFD diet supplemented with either white carrot powder (a food matrix control lacking carotenoids) or orange carrot powder (with carotenoids), while LFD (10% kcal from fat) was supplied to the negative control group.

Since carrots are one of the main sources of α-carotene and β-carotene (57), we first used HPLC to determine their concentration. Carotenoids were undetectable in LFD and HFD pellets, including HFD+WC. However, quantifiable amounts of α-carotene (0.25 ± 0.01 μmol/g) and β-carotene (0.18 ± 0.01 μmol/g) were detected in HFD+OC, highlighting the difference in carotenoid content due to the use of orange carrots. These findings align with documented evidence that α-carotene is one of the most abundant carotenoid in orange carrot cultivars (58). White carrots served as a control group for carotenoids and were added into the dietary pellets at the same ratio as the orange carrots. Thus, the only significant difference between the white and orange carrot pellets was their carotenoid content. The LFD, HFD, and HFD+WC groups lacked α-carotene or β-carotene in their circulation due to a lack of those compounds in their diet (Table 1). The HFD+OC group had serum levels of α-carotene (0.25 ± 0.07 μM) and β-carotene (0.26 ± 0.06 μM), which fall within the range of typical carotene concentrations in human serum (0.04-2.26 μM) including when taking oral β-carotene supplements or consuming a β-carotene-rich diet (59,60), though retinol levels were consistent. Colon tissues in the LFD, HFD, and HFD+WC groups lacked carotenoids. However, the HFD+OC group had α-carotene (0.72 ± 0.28 nmol/g) and β-carotene (0.43 ± 0.14 nmol/g), whereas retinol was undetectable in all groups. A significant portion of carotenoids are poorly absorbed and enter the colon (61), explaining the larger detected concentration in the colon compared to serum.

From week 2, all high-fat diet (HFD) groups had significantly higher body weights than the low-fat diet (LFD) group. Between Weeks 9 and 14, the HFD groups supplemented with white carrot (WC) and orange carrot (OC) exhibited higher body weights than the HFD alone group; however, by Week 20, only the HFD+WC group remained significantly greater. The HFD+OC group eventually gained the same body weight as the HFD and HFD+WC groups, demonstrating that these groups had similar weight results. In a previous study from our group, we found that orange carrot supplementation of the same dosage (20% w/w) in a HFD consisting of 60% kcals from fat over 15 weeks led to significant inhibition of body weight gain compared to the HFD group (32). Additionally, the white carrot supplementation in that study caused reduced body weight gain compared to the HFD control, though this was statistically insignificant. Diets classified as high-fat contain 20% to 60% fat and can be formulated with plant oils or animal fats like lard and beef tallow (62). Depending on the desired degree of metabolic disruption and its evolution, one can choose between diets with 45% or 60% fat. Diets with a high-fat content (≥60% energy) are less likely to be physiologically relevant (62). Therefore, in the current study, we chose to utilize a 45% fat diet, while in our pilot study, we used a 60% fat diet. The body weight results seen in the preliminary might not have been reflected in this study due to the difference in the dietary fat in the HFD pellets, as the fat-soluble carotenoids are most likely to be absorbed better under the 60% fat conditions.

EchoMRI analysis indicated significant increases in fat mass in all HFD groups compared to the LFD group, demonstrating the effects of high-fat diets without carrot supplementation. Notably, the HFD coupled with OC resulted in the highest lean mass among the groups. Given previous studies showing that carotenoids and polyphenols are inversely linked with sarcopenic symptoms, it is biologically feasible that these nutrients may protect skeletal muscle and function during calorie restriction in overweight and obese individuals (63). The findings of our study could confirm this concept, as the HFD supplemented with orange carrot resulted in the highest lean mass among the groups despite the general increase in fat mass shown in all HFD groups, which suggests that dietary carotenoids, like those found in orange carrots, may help maintain muscle mass and function even following a high-fat diet.

Obesity is characterized by low-grade chronic inflammation, as adipose tissue, an endocrine organ, secretes cytokines such as IL-6 and TNF-α (64). Although orange carrot supplementation did not inhibit weight gain, the carotenoids present decreased IL-6 and calprotectin levels, as the HFD+OC group had the most pronounced decrease in fecal calprotectin levels and the lowest colonic IL-6 concentrations, indicating a protective and anti-inflammatory effect of orange carrot supplementation in high-fat diet-induced obesity. The LFD group exhibited considerably lower serum levels of lipopolysaccharide (LPS), C-reactive protein (CRP), and lipopolysaccharide-binding protein (LBP) than the other groups, with no suppression of these markers detected in any of the carrot treatment groups.

Functional annotation and fold chance induction of key proteins that were involved in pathways like inflammation response, antioxidant activity, lipid metabolism, retinoid metabolism, and epithelial barrier management were portrayed in the GOnet plots that compared HFD+OC expression to HFD (**Figure 4**) and to HFD+WC (**Supplementary Figure 4**). These two comparisons were highlighted to investigate how the orange carrot led to changes in the distal colonic tissue under the challenge of the high-fat diet and, more importantly, to the interests of our lab, to emphasize how the carotenoid-rich dietary intervention induced further or differential changes in comparison to the carotenoid-deficient white carrots. In multiple cases, the white and orange carrot treatment groups induced identical changes compared to HFD, with no significant differences between the two carrot groups. Additionally, there were several instances where the HFD+WC induced higher or lower fold change induction in protein abundance compared to HFD to a greater extent than what was achieved by HFD+OC. This shows that components in the food matrix of the white carrots were able to induce a proteomic response under the high-fat diet challenge. To our surprise, the reduction of inflammatory markers in the colon was not a major mechanism induced by carrot supplementation under these high-fat conditions as abundances of key players in the microbial/pathogen associated molecular pattern response (TLR2, MYD88, NFKB1, IKBKB, TAB1, IL1RN) (38,41) typically influenced under a high-fat diet all remained unaffected. While TRAF6 and JAK2/STAT3 abundances were decreased by both HFD+WC and HFD+OC, this was not seen with JNK/p38 MAPK proteins as these partners were actually increased by HFD+WC, though unaffected by HFD+OC, and ERK1 (not ERK2) protein was elevated by both HFD+WC and HFD+OC. Stimulation of antioxidant activity in the colon was also not induced by the carrot diets of either color as proteins like TRX1, PRDX1, SOD1, CAT, KEAP1, GPX, NQO1, NOS1 were not influenced by either HFD+WC or HFD+OC; though the activity of SOD2 and PRDX2 were upregulated by white carrots alone or both carrot groups. Finally, proteins involved in intestinal fat regulation were indeed influenced by the carrot feeding, namely fatty acid oxidation facilitators like CPT1A, CPT2, ACOX1, and ACOX3, though these were relative to the LFD group. However, facilitators of intestinal fat metabolism like PPARα and PPARγ (46) were surprisingly not identified in our samples. Lipogenesis-related protein FASN was notably upregulated by both HFD+WC and HFD+OC relative to HFD, which has a protective effect on the intestinal barrier by promoting colonic MUC2 secretion via palmitoylation (47). The difference in proteome regulation between the carrot groups was apparent in epithelial barrier maintenance, as HFD+OC upregulates proteins in this category. Both DRG1, which has been seen to have a protective effect on epithelial junction proteins under the conditions of necrotizing enterocolitis (65) and MUC2 were uniquely upregulated by HFD+OC.

Taking a closer look into regulation induced by the supplementation of the carotenoid-rich orange carrots, proteomic analysis of the distal colon revealed that the relative abundances of several proteins involved in maintaining the mucosal barrier were enhanced by HFD+OC feeding. The STRING protein-protein interaction network showed how these targets are associated, with MUC2 as a major focal point of interconnectivity. Multiple key mucus-associated proteins, including MUC2, FCGBP, CLCA1, ZG16, AGR2, and KLK1, have been highlighted in another study involving the quantitative proteomic analysis of mucin granules from human colonic goblet-like LS174T cells (66). At the colonic level, the intestinal barrier is coated with a protective bilayer of mucus, of which the inner (glycocalyx) layer is adherent to the epithelial cells and impenetrable to bacteria, while the outer is loosely packed and colonized by microbes (67). MUC2, the key gel-forming secretory glycoprotein produced by intestinal goblet cells, is the predominant component of these mucus layers and improves barrier integrity by generating a skeletal network via cross-linkages with each other (68). Within the rough endoplasmic reticulum, AGR2 facilitates the assembly of MUC2 dimers via cysteine disulfide bonds, which are then shuttled to the Golgi apparatus for oxidative (O)-glycosylation and packaging into secretory granules (54). FCGBP is another abundant element of the mucus layer due to noncovalent binding with MUC2 (69). Furthermore, FCGBP is involved in the goblet cell immune response by binding to immunoglobulin G and can heterodimerize with trefoil factor 3 (TFF3) to impede microbial motility towards the barrier (53,70). Renewal of the mucus layer can be facilitated by CLCA1 through the cellular influx of calcium and efflux of bicarbonate (52,53), which shuttles the mucus from the inner layer towards the outer layer so that mucus can be freshly secreted from the goblet cells. The HFD+OC group in this study significantly promoted the abundance of MUC2 in the distal colon compared to all other groups, as shown by the proteomics and tissue staining. Additionally, FCGBP was elevated by HFD+OC to a level higher than HFD and HFD+WC groups near significantly as well as increased CLCA1 expression significantly more than HFD+WC. However, there was no significant effect on AGR2, and TFF3 was not identified by our proteomic analysis. Some intestinal components can directly interact with gut microbiota, either inflicting bactericidal effects or preventing microbes from readily passing through the mucosal layer. For example, ZG16 can bind to the peptidoglycan component of the cell wall in gram-positive bacteria, leading to bacterial aggregation that impairs motility towards the epithelial barrier (53); however, our dietary intervention did not cause any significant changes in ZG16 abundance. Although the function of KLK1 has not yet been specifically elucidated, its activity has been linked to these other mucus-associated proteins (66), which was further supported by this study as HFD+OC induced a significantly higher abundance of this protein than LFD and HFD as well as near significantly compared to HFD+WC. Additional mucins like MUC3, MUC4, MUC13, and MCAM were recognized via proteomics, all of which are membrane-bound mucins and not secretory proteins like MUC2 (51). The dietary intervention did not induce differences of statistical significance in these mucins, which may explain the lack of apparent changes in Alcian Blue staining in the colon sections. Stimulation of mucus secretion can be influenced by inflammatory pathways (e.g., NF-κB, MAPK signaling) (68), which may be why we did not see significant changes regarding these inflammatory proteins illustrated in the GOnet maps, suggesting that HFD+OC may have inhibited the expression of these proteins to a certain extent in an anti-inflammatory aspect but also promoted expression to facilitate mucus production. Unexpectedly, tight junction proteins CLDN3, CLDN7, TJP1, and TJP2 were not substantially promoted by the HFD+OC group. As previously mentioned, our group found that β-carotene treatment (10 nM-10 µM) promoted CLDN1 and OCLN protein expression in HT-29 cells (56); however, these particular tight junction proteins were not identified in this proteomics analysis. The upregulated abundance of MPTX1, MPTX2, and CCL6 by HFD+OC may indicate stimulated intestinal Paneth cell-mediated production of antimicrobial peptides and immune cell recruitment (42,55).

Several limitations exist in our study, some of which relate to the model of obesity. Firstly, there was no significant inhibition of HFD-induced body weight gain by white carrot and orange carrot feeding like what was seen during our prior study; in fact, there was a significant increase of body weight and fat mass in the HFD+WC fed mice compared to the HFD group. Previous research has shown that white carrots have a high sugar content but lower levels of ascorbic acid, antioxidants, and tetraterpenoids than other colored carrots (71). This increased sugar content could likely contribute to the increased weight observed in the white carrot-fed group (HFD+WC). However, in our previous investigation, a high-fat model of 60% kcals from fat may have led to greater absorption of the fat-soluble carotenoids in the orange carrots due to the higher dietary lipid content and caused a more pronounced anti-obesogenic effect. While there may not be overall changes in body weight at 45% HFD, tissue-specific regulation could still be happening in other organs like the colon, as demonstrated by this study, in addition to sites relevant to both obesity and carotenoid bioactivity such as the liver and adipose tissue that can be investigated in our future experiments. Secondly, the dietary intervention of carrots did not lead to significant decreases in circulatory LPS, LBP, and CRP, which indicated that there was no effect on metabolic endotoxemia, at least with these markers. Regarding the quantitative proteomics of the distal colon tissue, the HFD versus LFD volcano plot did not show any significant upregulation or downregulation in this particular proteome comparison, which is curious since there were significant differences in body weight gain and serum inflammatory markers when comparing these control groups over the 20-week period. This lack of a proteomic response could indicate that pathways like obesity-associated colon-specific low-grade inflammation and barrier impairment were not induced in this model despite similar studies achieving gut dysfunction with 45% kcals from fat in a similar time frame (72,73). Furthermore, key carotenoid metabolism proteins like BCO1 and BCO2 cleavage enzymes and retinoid conversion enzyme lecithin: retinol acyltransferase (LRAT) (5) were not identified in our samples. Of those recognized, retinoid metabolism and receptors were not significantly regulated by HFD+OC in the colon tissue (ALDHs, RDHs, DHRSs, and RXRB), which could indicate that the large intestine is not a major site of carotenoid conversion to vitamin A and has instead been done in the small intestine. However, SR-B1 (Scarb1) was identified in the proteomics, indicating that carotenoids can be taken up into the colon. Therefore, retinoid signaling may be limited in the colon under these study design conditions (i.e., 45% kcals from fat). Some additional key mucus-associated proteins were not identified in our samples, such as TFF3, ly6/plaur domain-containing protein 8 (LYPD8), and resistin-like molecule beta (RELMꞵ). Besides heterodimerization with FCGBP, TFF3 can improve barrier integrity by promoting the E-cadherin/β-catenin connections at cell-cell junctions and reducing pore-forming claudin-2 expression (74). LYPD8 and RELMꞵ both target members of the gut microbial community as LYPD8 can inhibit motility of flagellated bacteria (75) and RELMβ, which is synthesized by distal colonic goblet cells, can perforate the cell membranes of gram-negative bacteria (76); RELMꞵ was identified in our analysis, but not in every experimental replicate. Additionally, proteins investigated in our HT-29 study (56) were not identified in the proteomics, such as TLR4, CD14, NF-κB p65, CLDN1, and OCLN. Lastly, our data visualization tool of GOnet required the use of gene abbreviations in lowercase form, which may not match common protein nomenclature (e.g., Scar-b1 versus SR-B1), as our raw proteomic data generated from the Proteome Discoverer was supplied with gene names, though UniProt IDs have been provided.

In conclusion, our study demonstrated that supplementation with orange carrots as a source of carotenoids in a high-fat diet significantly upregulates colonic mucus synthesis and secretion. This enhancement of the gut barrier integrity is a promising indicator of the potential protective effects against diet-induced gut dysfunction and inflammation. These findings underscore the therapeutic potential of carotenoids in managing and preventing gastrointestinal disorders associated with high-fat diets. In other studies, β-carotene (60 mg/kg in diet for 28 days) increased ileal mRNA levels of *Muc2, Zo-1, Zo-2*, and *Ocln* in Hyline Brown chicks (77). Additionally, β-carotene (50 mg/kg BW for two weeks) increased colonic mRNA levels of *Muc2* and *Muc3* under vitamin A-deficient conditions (78). As for inflammation, β-carotene supplementation at 40 and 80 mg/kg body weight (BW)/day for 26 days was proven to increase growth performance intestinal morphology and decrease inflammation in weaning piglets via decreased NF-κB and proinflammatory cytokine signaling (79). Since the animals in these studies were treated with pure carotenoids and conducted under varying models, the findings from this study provided unique evidence that α-carotene and β-carotene supplementation from a whole food source promoted colonic mucus-associated protein activity under high-fat diet conditions.

Expanding on the substantial elevation of mucus-associated proteins reported after orange carrot supplementation, future studies could investigate the gut microbiome, specifically regarding mucus-interacting gut bacteria such as *Akkermansia muciniphila*. Other carotenoid supplementation studies found that carotenoids have the prebiotic capability of promoting *Bifidobacterium, Lactobacillus,* and *Akkermansia muciniphila* abundance (1,61,80). On a related note, extensive research has been done connecting dietary polyphenols to mucin-2 and *Akkermansia muciniphila,* including under HFD conditions (81,82). MUC2 has also been shown to enhance the binding of secretory immunoglobulin A (sIgA) and antimicrobial peptides to the gut epithelium, an essential interaction in preserving gut barrier integrity and immune defense. Furthermore, earlier studies have indicated that short-chain fatty acids (SCFAs) are crucial in regulating mucin production, emphasizing the associated roles of dietary components and gut microbiota in regulating mucosal immunity (83,84). Therefore, we hypothesize that the high-fat diet supplemented with orange carrots (HFD+OC) in this study led to a notable increase in the populations of beneficial SCFA-producing and mucus-interacting microbes, like *Bifidobacterium*, *Lactobacillus*, and *Akkermansia muciniphila* compared to other dietary intervention groups. Understanding these relationships may provide more information on how carotenoids influence gut health and host metabolism.

Future studies will incorporate various different colored carrot powders, composed of different carotenoid profiles, to address whether other major dietary carotenoids can mitigate the effects of diet-induced obesity. To investigate the impacts of various carotenoids, we are now employing a model that comprises a low-fat diet, high-fat diet, HFD with white carrots, HFD with orange carrots, HFD with yellow carrots, and HFD with red carrots as a prospective future approach. Furthermore, liver and adipose tissues, as well as the gut microbiome, will be investigated to reveal the role of dietary carotenoids in hots-microbiota interactions in obesity.

## Data Availability Statement

Data is contained within the article or supplementary material. The mass spectrometry proteomics data have been deposited to the ProteomeXchange Consortium (https://proteomecentral.proteomexchange.org/) (85) via the PRIDE partner repository (86, 87) with the data set identifier **PXD054150**.

## Acknowledgments

This material is based on work that is supported by the National Institute of Food and Agriculture, U.S. Department of Agriculture, under award number 2022-67018-37188. We are grateful to other members of the Eroglu lab who were involved in this work at North Carolina State University, Ms. Ezgi Ersoy and Ms. Chloe Cable. We also thank the North Carolina Food Innovation Lab (NCFIL) and Research Diets, Inc., for generating carrot powders and incorporating them into intervention arms, respectively. We are very appreciative to Glicerio Ignacio, DVM, DACAM (The McConnell Group, Kannapolis, NC, as he provided tremendously to our study with respect to work in the vivarium with animal husbandry, blood draws, and necropsy. The proteomic LC-MS/MS analysis could not have been done without the hard work of the members of the Molecular Education, Technology, and Research Innovation Center (METRIC) at NC State University. Finally, the Alcian Blue and MUC2 staining of the distal colon sections were conducted by NC State Anatomical Pathology Histology Lab at the College of Veterinary Medicine.

## Conflicts of Interest

The authors declare no conflicts of interest. The funders had no role in the study’s design, in the collection, analyses, or interpretation of data, in the writing of the manuscript, or in the decision to publish the results.

## Institutional Review Board Statement

The study was approved by the Institutional Animal Care and Use Committee (IACUC) at North Carolina State University (protocol code #21-019 and date of approval: 10/04/2021).

## Abbreviations

ACOX1: Acyl-Coenzyme A oxidase 1
AGR2: Anterior gradient protein 2 homolog
ALDH1A2: Retinal dehydrogenase 2
ALDH1A3: Aldehyde dehydrogenase 3
BCO1: ꞵ-carotene 15,15’-dioxygenase
CAMKK2: Calcium/calmodulin-dependent protein kinase kinase 2
CAT: Catalase
CD36: Cluster determinant 36
CLCA1: Calcium-activated chloride channel regulator 1
CLDN1: Claudin-1
CLDN3: Claudin-3
CLDN7: Claudin-7
CPT1A: Carnitine O-palmitoyltransferase
CPT2: Choline/carnitine acyltransferase domain-containing protein
CRP: C-reactive protein
DHRS: Dehydrogenase/reductase SDR
DRG1: Developmentally-regulated GTP-binding protein 1
ERK2: Extracellular signal-regulated kinase 1
FABP2: Intestinal fatty acid binding protein
FASN: Fatty acid synthase
FCGBP: Iggfc-binding protein
Fut2: Galactoside alpha-(1,2)-fucosyltransferase 2
GPX1: Glutathione peroxidase 1
HFD: High fat diet
HT-29: Human adenocarcinoma colon epithelial cells
IBD: Inflammatory bowel disease
IKBKB/IKK-ꞵ: Inhibitor of kinase NF-κb subunit beta
IL-6: Interleukin 6
IL1RN: Interleukin 1 receptor antagonist
ISX: Intestine-specific homeobox
JAK2: Tyrosine-protein Janus Kinase 2
JNK: Jun N-terminal kinase
KEAP1: Kelch-like ECH associated protein
KEGG: Kyoto Encyclopedia of Genes and Genomes.
KLK1: Kallikrein-1
LBP: Lipopolysaccharide-binding protein
LC-MS/MS: Liquid Chromatography with tandem mass spectrometry
LFD: Low fat diet
LPS: Lipopolysaccharide
MAPK: Mitogen-activated protein kinase
MCAM: Mucin 18
MPTX: Mucosal pentraxin
MUC-2: Mucin-2
MUC13: Mucin 13
MUC3: Mucin 3
MUC4: Mucin 4
MYD88: Myeloid differentiation primary response 88
NQO1: NAD(P)H quinone dehydrogenase 1
NF-κb: Nuclear factor kappa-light-chain-enhancer of activated B cells
NFKB1: NF-κb p105 subunit
NOS1: Nitric oxide synthase
OC: Orange carrot
OCLN: Occludin
PRDX1: Peroxiredoxin-1
RDH: Retinol dehydrogenase
RXRꞵ: Retinoid X receptor beta
SCD2: Stearoyl-Coenzyme A desaturase 2
SOD1: Superoxide dismutase
SR-B1: Scavenger receptor class B type 1
STAT3: Signal transducer and activator of transcription 3
TAB1: Transforming growth factor- β-activated kinase 1 regulatory subunit
TJP1: Zonula occludens-1
TJP2: Zonula occludens-2
TLR: Toll-like receptor
TLR4: Toll-like receptor 4
Tmtpro: Tandem Mass Tag pro
TRX1: Thioredoxin
WC: White carrot
ZG16: Zymogen granule membrane protein 16

## Notes

### Competing Interest Statement

The authors have declared no competing interest.

