## Supplemental for "The Biochemical Effects of Carotenoids in Orange Carrots on the Colonic Proteome in a Mouse Model of Diet-induced Obesity"

### Supplementary Materials

**A)**

| | D12450HB (LFD) | D12451B (HFD) | White Carrot (HFD+WC) | Orange Carrot (HFD+OC) |
|---|---|---|---|---|
| Ingredient | gm | gm | gm | gm |
| Casein | 200 | 200 | 186.7 | 186.7 |
| L-Cystine | 3 | 3 | 3 | 3 |
| Corn Starch | 452.2 | 72.8 | 0 | 0 |
| Maltodextrin 10 | 75 | 100 | 72.7 | 72.7 |
| Sucrose | 172.8 | 172.8 | 172.8 | 172.8 |
| Cellulose, BW200 | 50 | 50 | 9.1 | 9.1 |
| Soybean Oil | 25 | 25 | 25 | 25 |
| Lard | 20 | 177.5 | 177.5 | 177.5 |
| Mineral Mix, S10026 | 10 | 10 | 10 | 10 |
| Dicalcium Phosphate | 13 | 13 | 13 | 13 |
| Calcium Carbonate | 5.5 | 5.5 | 5.5 | 5.5 |
| Potassium Citrate, 1 H2O | 16.5 | 16.5 | 16.5 | 16.5 |
| **Vitamin Mix, V10001** | 10 | 10 | 10 | 10 |
| Choline Bitartrate | 2 | 2 | 2 | 2 |
| FD&C Blue Dye #1 | 0.05 | 0.05 | 0.05 | 0.05 |
| **White carrot powder** | **0** | **0** | **175.6** | **0** |
| **Orange carrot powder** | **0** | **0** | **0** | **175.6** |
| **Total** | **1055.05** | **858.15** | **879.45** | **879.45** |
| gm | | | | |
| Protein | 177.0 | 177.0 | 177.0 | 177.0 |
| Carbohydrate | 710.0 | 355.6 | 355.6 | 355.6 |
| Fat | 45.0 | 202.5 | 202.5 | 202.5 |
| Fiber | 50.0 | 50.0 | 50.0 | 50.0 |
| gm% | | | | |
| Protein | 16.8 | 20.6 | 20.1 | 20.1 |
| Carbohydrate | 67.3 | 41.4 | 40.4 | 40.4 |
| Fat | 4.3 | 23.6 | 23.0 | 23.0 |
| **Carrot Powder** | **0.0** | **0.0** | **20.0** | **20.0** |
| kcal | | | | |
| Protein | 708 | 708 | 708 | 708 |
| Carbohydrate | 2840 | 1422.4 | 1422 | 1422 |
| Fat | 405 | 1822.5 | 1823 | 1823 |
| Total | 3953 | 3953 | 3953 | 3953 |
| kcal% | | | | |
| Protein | 18 | 18 | 18 | 18 |
| Carbohydrate | 72 | 36 | 36 | 36 |
| Fat | 10 | 46 | 46 | 46 |
| kcal/gm | 3.75 | 4.61 | 4.49 | 4.49 |

**B)**

| | | |
|---|---|---|
| **Carbohydrate** | Sucrose, Fine Granulated | 78.42 g |
| **Vitamin** | Vitamin E Acetate, 50% | 10.00 g |
| **Vitamin** | Niacin (a.k.a. B3, Nicotinic Acid) | 3.00 g |
| **Vitamin** | Biotin, 1% | 2.00 g |
| **Vitamin** | Pantothenic Acid, d, Calcium (a.k.a. B5) | 1.60 g |
| **Vitamin** | Vitamin D3, 100,000 IU/gm | 1.00 g |
| **Vitamin** | Vitamin B12, Cyanocobalamin, 0.1% | 1.00 g |
| **Vitamin** | Vitamin A Acetate, 500,000 IU/gm | 0.80 g |
| **Vitamin** | Pyridoxine HCl (a.k.a. B6) | 0.70 g |
| **Vitamin** | Riboflavin (A.K.A. B2) | 0.60 g |
| **Vitamin** | Thiamine HCl (a.k.a. B1) | 0.60 g |
| **Vitamin** | Folic Acid | 0.20 g |
| **Vitamin** | Menadione Sodium Bisulfite | 0.08 g |
| | Total: | 100.00 g |

**Supplementary Table 1: Ingredients of experimental diet pellets.** Compositional breakdown of **A)** dietary pellets and **B)** vitamin (V10001) mix formulated by Research Diets, Inc.

| Fraction No. | ACN (%) | ACN (μL) | TEA (0.1%) (μL) |
|---|---|---|---|
| Wash | 5.0 | 50 | 950 |
| 1 | 10.0 | 100 | 900 |
| 2 | 12.5 | 125 | 875 |
| 3 | 15.0 | 150 | 850 |
| 4 | 17.5 | 175 | 825 |
| 5 | 20.0 | 200 | 800 |
| 6 | 22.5 | 225 | 775 |
| 7 | 25.0 | 250 | 750 |
| 8 | 50.0 | 500 | 500 |

**Supplementary Table 2: Fractionation of TMT-labeled peptides.** Volume of LC-MS grade acetonitrile (ACN) and triethylamine (TEA) to fractionate TMT-labeled peptides. Fraction 8 was chosen for the proteomic LC-MS/MS analysis.

| | | -2 in % | | -1 in % | | M+ in % | +1 in % | | +2 in % | |
|---|---|---|---|---|---|---|---|---|---|---|
| **Mass Tag** | **Reporter Ion Mass** | **-2x13C** | **-13C-15N** | **-13C** | **-15N** | **-** | **+15N** | **+13C** | **+15N+13C** | **+2x13C** |
| 126 | 126.127726 | 0 | 0 | 0 | 0 | 100% | 0.31 | 9.09 | 0.02 | 0.32 |
| 127N | 127.124761 | 0 | 0 | 0 | 0.57 | 100% | 0 | 9.79 | 0 | 0.33 |
| 127C | 127.131081 | 0 | 0 | 0.84 | 0 | 100% | 0.23 | 8.4 | 0.02 | 0.27 |
| 128N | 128.128116 | 0 | 0 | 0.82 | 0.65 | 100% | 0 | 8.13 | 0 | 0.26 |
| 128C | 128.134436 | 0 | 0 | 1.44 | 0 | 100% | 0.34 | 6.26 | 0 | 0.17 |
| 129N | 129.131471 | 0 | 0.14 | 1.3 | 0.89 | 100% | 0 | 7.52 | 0 | 0.12 |
| 129C | 129.13779 | 0.13 | 0 | 2.59 | 0 | 100% | 0.32 | 6.07 | 0.01 | 0.09 |
| 130N | 130.134825 | 0.13 | 0 | 2.41 | 0.27 | 100% | 0 | 5.58 | 0 | 0.1 |
| 130C | 130.141145 | 0.25 | 0 | 3.22 | 0 | 100% | 0.28 | 5.06 | 0 | 0.06 |
| 131N | 131.13818 | 0.03 | 0 | 2.78 | 0.63 | 100% | 0 | 4.57 | 0 | 0.12 |
| 131C | 131.144499 | 0.08 | 0 | 3.94 | 0 | 100% | 0.45 | 3.33 | 0 | 0.02 |
| 132N | 132.141535 | 0.07 | 0.01 | 3.14 | 0.73 | 100% | 0 | 3.4 | 0 | 0.03 |
| 132C | 132.147855 | 0.08 | 0 | 3.65 | 0 | 100% | 0.5 | 1.97 | 0 | 0 |
| 133N | 133.14489 | 0.15 | 0.01 | 3.58 | 0.72 | 100% | 0 | 1.8 | 0 | 0 |
| 133C | 133.15121 | 0.18 | 0 | 4.14 | 0 | 100% | 0.4 | 1.11 | 0 | 0 |
| 134N | 134.148245 | 0.3 | 0.03 | 5.49 | 0.62 | 100% | 0 | 1.14 | 0 | 0 |

**Supplementary Table 3: The quantification method used in PD** incorporated the parameters for reporter ions and isotope correction factors as defined by the product data sheet for TMTpro product number A44521, lot no. YG373065.

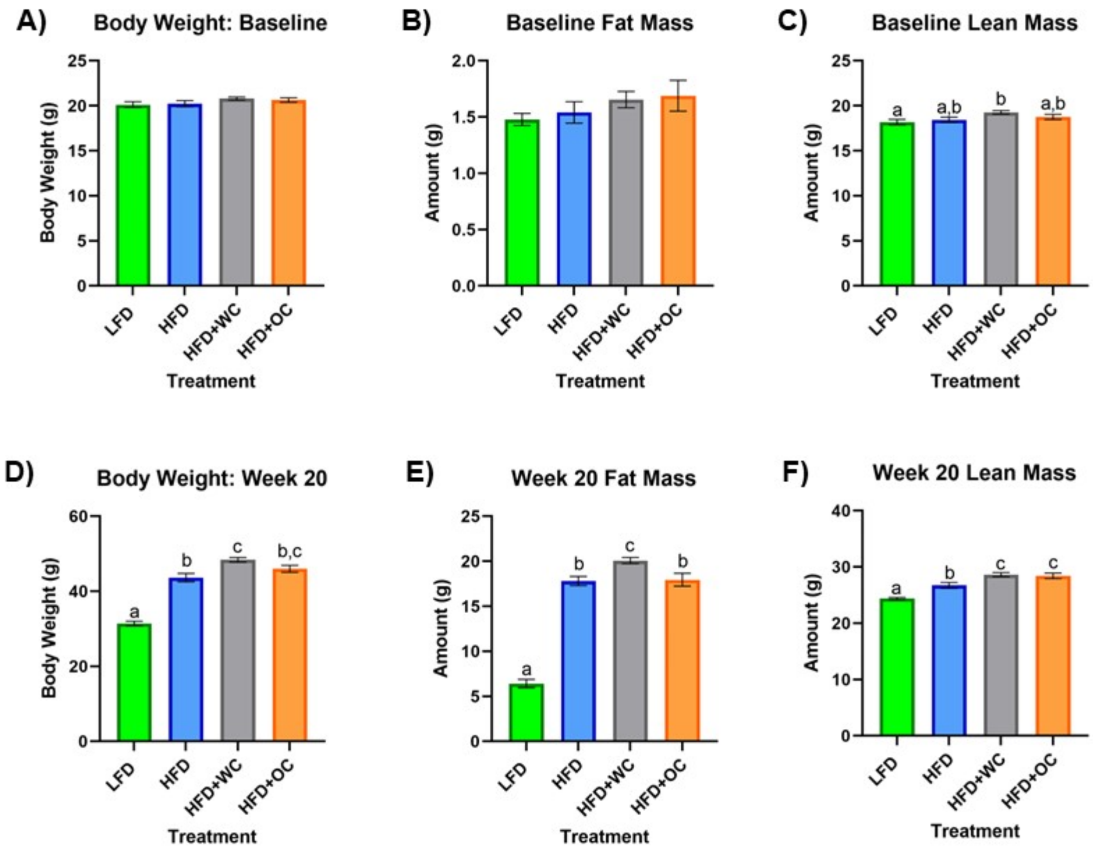

**Supplementary Figure 1: Individual measurements of body composition at baseline and endpoint**. **A-C)** Baseline measurements of **A)** body weight, **B)** fat mass, and **C)** lean mass at Week 0 before dietary intervention. **D-F)** Endpoint measurements of **D)** body weight, **E)** fat mass, and **F)** lean mass at Week 20. Fat mass (**B&E)** and lean mass (**C&F)** measurements determined by EchoMRI. LFD, HFD, HFD+WC, HFD+OC n=20. Values are means ± SEM. Body weight, fat mass, and lean mass measurements were analyzed by one-way ANOVA with post-hoc Tukey HSD. Letter differences indicate statistical significance ($p < 0.05$).

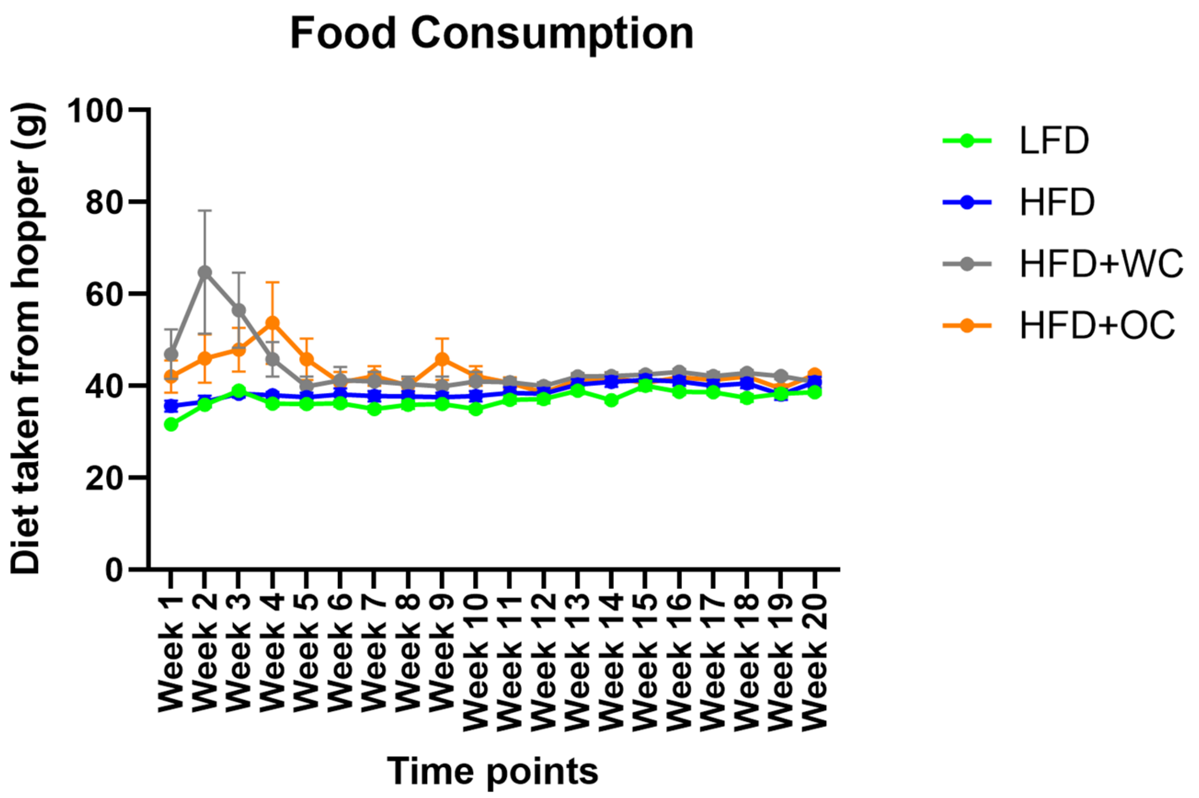

**Supplementary Figure 2: Weekly food consumption.** Measurements of diet pellets taken from the food hopper in the cages on a weekly basis. Weight of pellets in the hopper only (not including any food on the floor of the cage) was recorded at the time of administration and then the amount leftover when the week elapsed, reporting the difference between the two points. A two-way mixed ANOVA with post hoc Tukey HSD to illustrate the differences in food consumption over time.

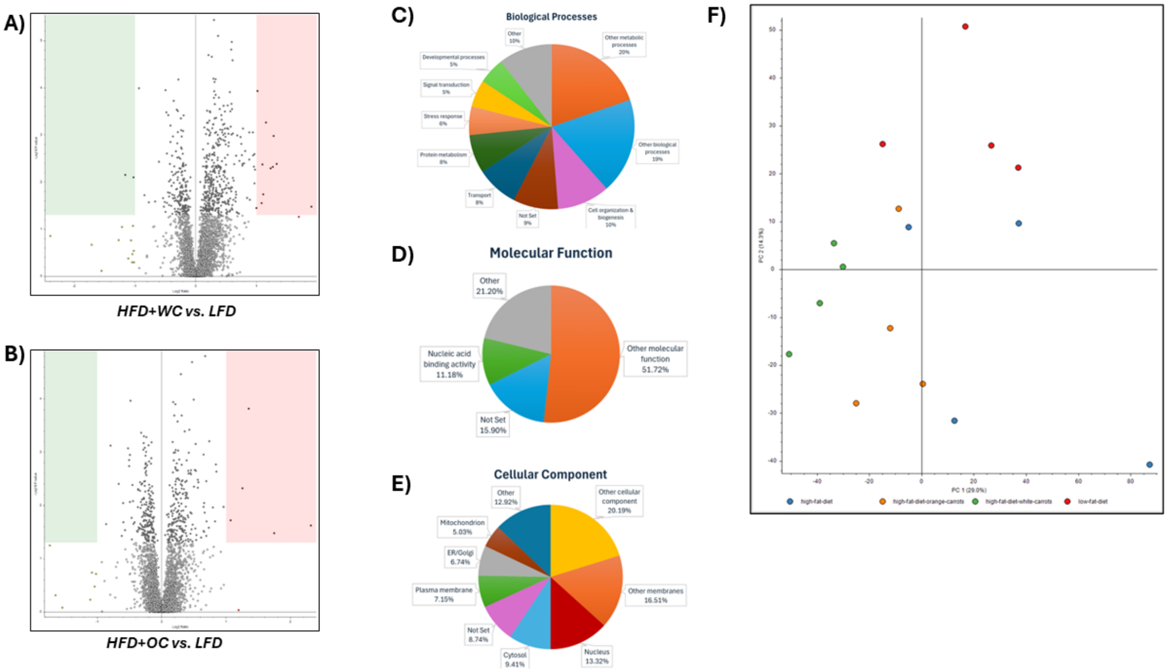

**Supplementary Figure 3: Additional comparisons from quantitative proteomics. A-B)** Volcano plots of differentially expressed peptides upon comparison of abundances between **A)** HFD+WC vs LFD and **B)** HFD+OC vs LFD. **C-E)** pie charts of identified protein KEGG characteristics, **C)** Biological Process, **D)** Molecular function, **E)** Cellular Component. **F)** PCA plot of identified peptides across treatment groups

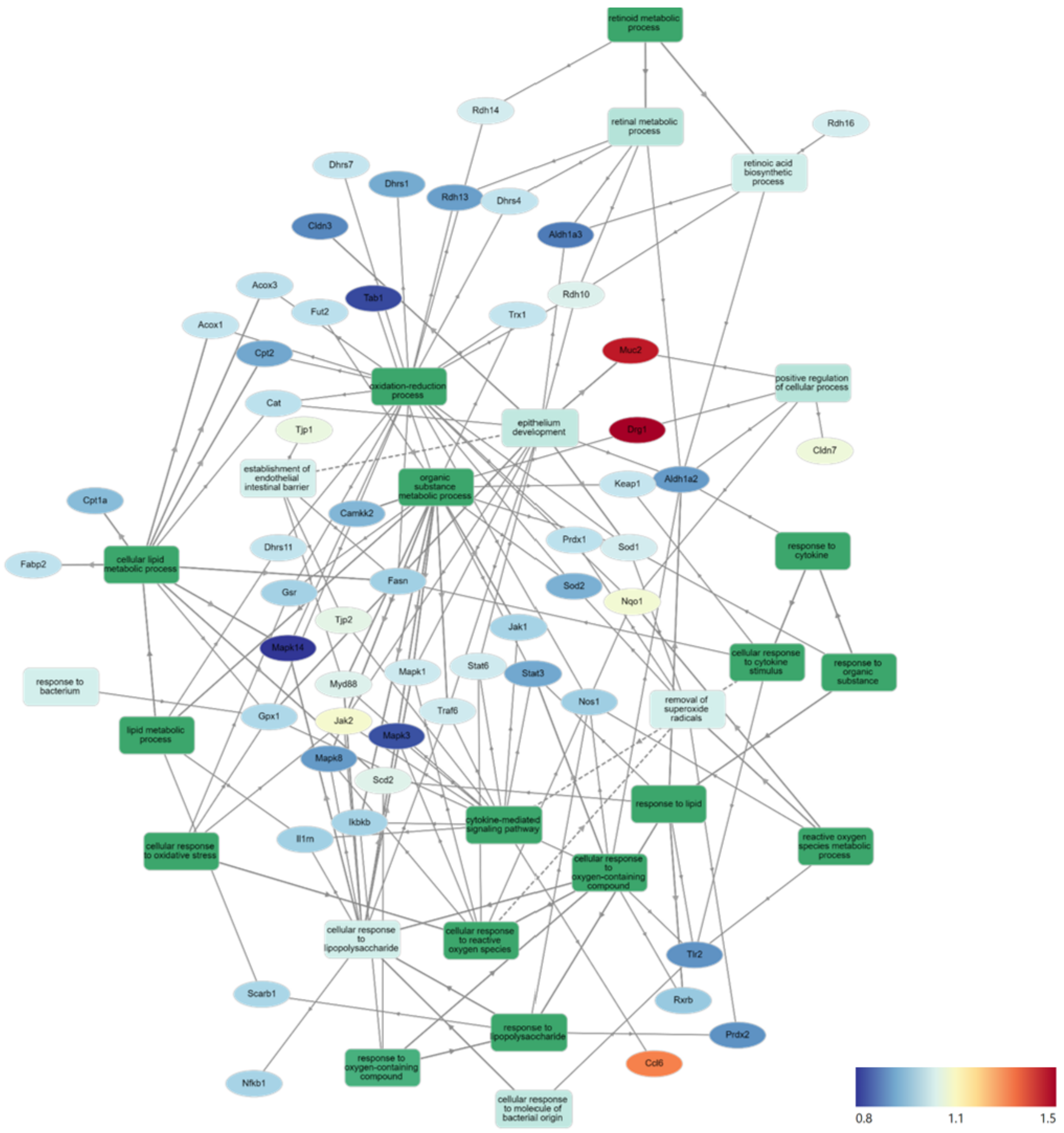

**Supplementary Figure 4: GOnet Map of HFD+OC/HFD+WC comparison.** Gene Ontology Network (GOnet) map of the same proteins depicted in Figure 4 with relative quantitative comparison between HFD+OC versus HFD+WC (color of protein nodes and legend correspond to HFD+OC/HFD+WC fold change). KEGG Pathway: Biological Process
